# Reconstructing biologically coherent cellular profiles from imaging-based spatial transcriptomics

**DOI:** 10.64898/2026.03.08.710395

**Authors:** Long Yuan, Youyun (Peter) Zheng, Shuming Zhang, Rameen Beroukhim, Atul Deshpande

**Affiliations:** Department of Immunology, Johns Hopkins University School of Medicine, Baltimore, MD, USA; Department of Computer Science, Johns Hopkins University Whiting School of Engineering, Baltimore, MD, USA; Department of Medical Oncology and Cancer Biology, Dana Farber Cancer Institute, Boston, MA, USA; Department of Biomedical Informatics, Harvard Medical School, Boston, MA, USA; Bloomberg-Kimmel Institute for Cancer Immunotherapy, Baltimore, MD, USA; Sidney Kimmel Comprehensive Cancer Center, Johns Hopkins University School of Medicine, Baltimore, MD, USA; Convergence Institute, Johns Hopkins University, Baltimore, MD, USA; Department of Cancer Biology, Dana-Farber Cancer Institute, Boston, Massachusetts, USA; Department of Medical Oncology, Dana-Farber Cancer Institute, Boston, Massachusetts, USA; Broad Institute of MIT and Harvard, Cambridge, MA, USA; Department of Medicine, Harvard Medical School, Boston, MA, USA; Department of Electrical and Computer Engineering, Johns Hopkins University, Baltimore, MD, USA; Data Science and AI Institute, Johns Hopkins University, Baltimore, MD, USA

**Keywords:** Spatial transcriptomics, 3D segmentation correction, segmentation diagnostics, cellular reconstruction

## Abstract

In imaging-based spatial transcriptomics, transcript-to-cell assignment shapes downstream biological interpretation including cell typing, ligand-receptor inference, and niche characterization. However, two-dimensional segmentation of volumetric tissue often yields mixed cellular profiles, while cells without detected nuclei are missed entirely, distorting the aforementioned downstream analyses. We present TRACER, which refines cellular representations in imaging-based transcriptomics by leveraging gene–gene coherence and spatial co-localization of transcripts observed directly in the data, without requiring external annotations or reference atlases. TRACER resolves mixed cellular profiles and reconstructs partial cells whose nuclei are not detected, enabling more complete representation of cells within the tissue section. We also introduce coherence-based metrics that quantify transcriptional purity and conflict, enabling platform-agnostic benchmarking of segmentation quality. Across diverse platforms, tissues, and segmentation methodologies, TRACER consistently and reproducibly improves the coherence of cellular profiles and the quality of downstream analyses.

## 1 Introduction

Imaging-based spatial transcriptomics enable the measurement of gene expression at subcellular resolution within the spatial context of the tissue, offering unprecedented opportunities to study tissue architecture, cell states, and cell–cell interactions in situ [1– 3]. Central to these analyses is the accurate assignment of individual transcripts to cells, the fundamental unit for downstream interpretation, including cell-type identification, differential expression analysis, and spatial niche modeling [4, 5]. Errors in imaging-based cell segmentation—typically used for transcript assignment—propagate directly into biological conclusions, making the accuracy of this step a critical determinant of analytical fidelity [4–7].

Despite rapid advances in imaging technologies and computational methods, accurate cell segmentation in spatial transcriptomics remains a challenge. Many pipelines rely on two-dimensional (2D) segmentation derived from nuclear staining, nuclear expansion heuristics, or membrane markers when available [4, 5, 8, 9]. Segmentation methods must contend with variable signal quality, densely packed tissues, and the intrinsic thickness of biological sections. Recent work has highlighted that tissue sectioning itself introduces ambiguity in cellular representation. Tissue sectioning causes contamination by the vertical environment of cells, specifically by partially captured cells with out-of-plane nuclei [10, 11]. Yet most approaches rely on 2D segmentation— derived from nuclear staining, nuclear expansion heuristics, or membrane markers when available—under the assumption that all transcripts within the 2D boundary belong to the same cell[4, 8, 9, 12]. As a result, segmented boundaries may not accurately represent cellular profiles, and subsequent analyses often exhibit mixed transcriptional signatures that are difficult to interpret [13, 14] (Fig. 1A).

**Figure 1.**
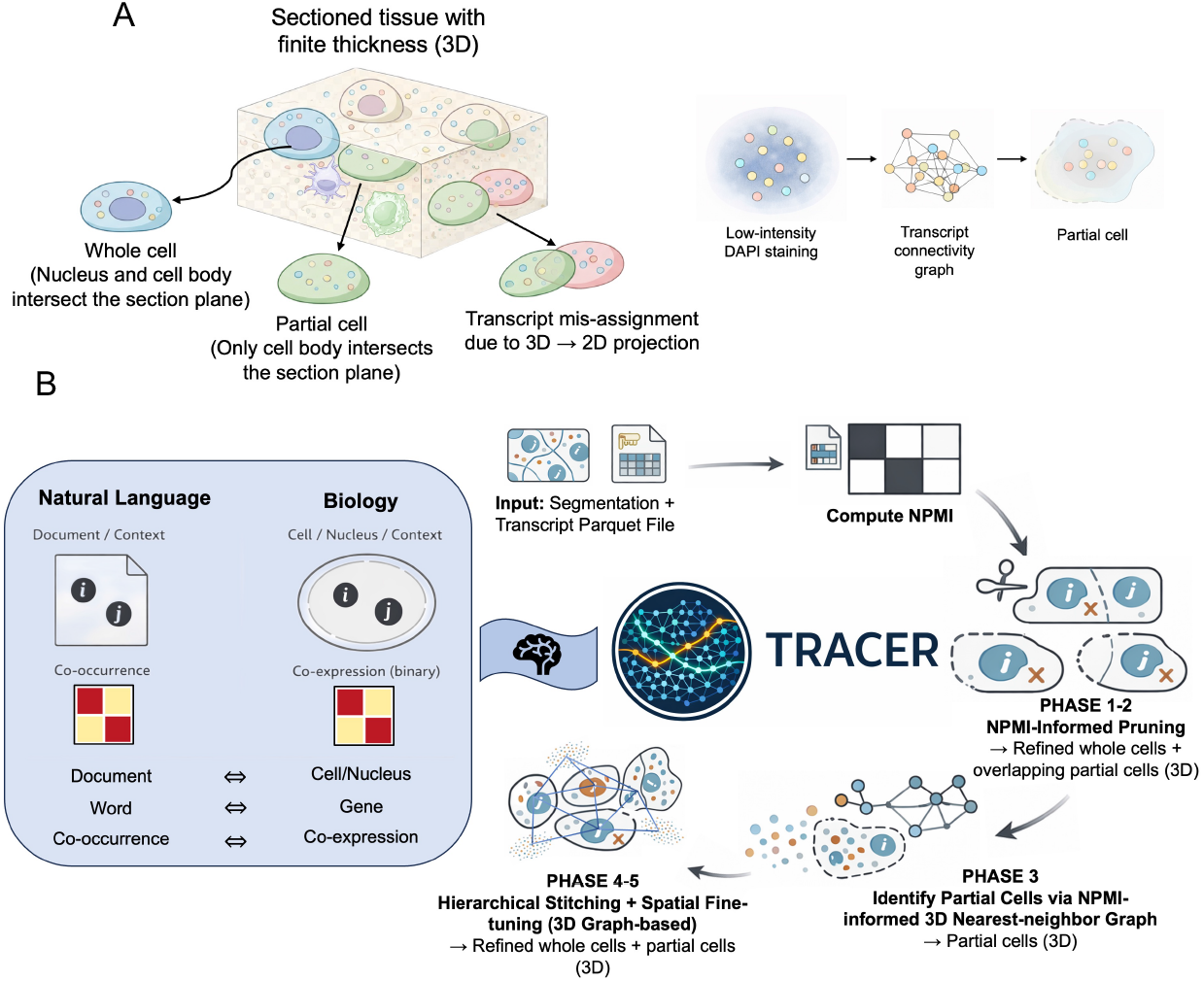
Biological motivation and algorithmic overview of TRACER. (A) 2D-segmentation on non-negligible tissue thickness produces errors in imaging-based spatial transcriptomics. Cells located above or below the imaging plane may intersect the section only partially, producing whole cells (nucleus and cytoplasm captured in-plane) and partial cells (cytoplasmic transcripts captured without an in-plane nucleus). Overlap of cellular profiles along the z-axis causes transcripts from different cells to project into the same two-dimensional footprint, resulting in systematic transcript misassignment. In addition, low-intensity or out-of-plane nuclei can leave clusters of transcripts unassigned under standard segmentation. These transcripts can be represented as a spatial transcript-connectivity graph, allowing coherent partial cellular profiles to be identified. (B) Conceptual framework and workflow of TRACER. TRACER draws an analogy to natural language processing in which nuclei define local context windows and genes are treated as binary events, enabling estimation of a data-driven gene–gene co-expression prior via normalized pointwise mutual information (NPMI). This prior guides: (1) NPMI-informed extraction of transcriptionally coherent whole cells and partial cells (Phases 1–2); (2) identification of partial cellular profiles from spatially structured unassigned transcripts using an NPMI-informed three-dimensional nearest-neighbor graph (Phase 3); and (3) hierarchical stitching and spatial fine-tuning of reconstructed cellular entities using a Δ*C* coherence-optimization criterion (Phases 4–5). The final output is a refined, three-dimensional–aware cellular representation that accounts for segmentation errors, incomplete cellular profiles, and transcript misassignment.

At the same time, there is growing recognition that imaging-based spatial transcriptomics data lack a clear gold standard for evaluating segmentation quality [15, 16]. Metrics developed for natural image segmentation focus on geometric overlap or boundary accuracy, which do not readily translate to biological context where cell boundaries are fuzzy, overlapping, and functionally defined [17]. Recent work using matrix factorization of local molecular neighborhoods [18] further demonstrated that segmentation-induced transcript admixture can dominate downstream analyses in imaging-based spatial transcriptomics, underscoring the need for approaches that explicitly address transcript mixing and segmentation uncertainty. As a result, segmentation methods are often assessed indirectly through downstream tasks, such as clustering or marker gene expression, without principled segmentation-agnostic diagnostics to quantify transcriptional coherence or mixing at the cellular level [4, 19].

Here, we introduce an information-theoretic approach for refining cellular representations and evaluating segmentation fidelity in spatial transcriptomics. We present TRACER (Tissue Reconstruction via Associative Clique Extraction and Relation-mapping), a computational framework designed to address the limitations of segmentation in imaging-based spatial transcriptomics by leveraging transcriptional associations directly from the data (Fig. 1B). TRACER uses gene–gene coherence and spatial colocalization to refine cellular representations without requiring external annotations, supervised training, or GPU acceleration. We also provide coherence-based metrics that enable evaluation of transcript-to-cell assignment across platforms and methodologies.

## 2 Results

### 2.1 TRACER uses transcriptional coherence to refine transcript assignment

TRACER is a computational framework for refining cellular representations in imaging-based spatial transcriptomics. TRACER leverages gene-gene associations and spatial proximity to address erroneous transcript assignments and to recover fragments of cells whose nuclei lie outside the section plane. The core design of TRACER is motivated by an analogy between natural language processing and spatial transcriptomics, treating cells (or nuclei) as contexts and gene presence as events, enabling robust estimation of gene–gene co-expression independent of transcript counts (Fig. 1B).

Using nuclear segmentation as a reliable context window, TRACER computes a gene–gene normalized pointwise mutual information (NPMI) matrix directly from the data [20–22]. These associations are then used to guide conservative transcript pruning, recovery of partial cell fragments, and constrained hierarchical stitching of fragmented cellular profiles into a three-dimensional (3D) representation (Fig. 1B). The stitching criterion, based on a coherence-change metric (Δ*C*), selectively merges spatially proximal fragments that are transcriptionally coherent while preventing incompatible merges; we validate this approach in a controlled 3D simulation (Supplementary Fig. 1 ; see Methods).

We distinguish between two stages of TRACER output used throughout this study. TRACER-stitched refers to the intermediate output obtained after NPMI-guided transcript pruning and Δ*C*-based hierarchical stitching, which yields transcriptionally coherent whole cells while preserving all spatially connected transcript components. This stage primarily resolves transcript mixing and hybrid cellular profiles arising from segmentation errors. TRACER-fine-tuned refers to a subsequent spatial refinement step in which each stitched entity is subjected to a spatial connectivity constraint. Specifically, disconnected transcript components that lie beyond a user-defined distance threshold from the dominant connected component are removed and reassigned through an additional constrained stitching pass. This step eliminates spatially inconsistent outliers while preserving the dominant transcriptional program, yielding the final refined set of whole cells, partial cells, and reconstructed partial cells used for downstream analyses (see Methods).

Unless otherwise stated, results are shown for both TRACER-stitched and TRACER-fine-tuned outputs to illustrate the impact of coherence-based transcript reassignment alone and in combination with spatial fine-tuning.

### 2.2 NPMI captures lineage-specific gene programs and enables quantitative purity and conflict scoring

We first evaluated whether the NPMI values computed from nuclear contexts captures biologically meaningful gene–gene relationships. In Xenium breast cancer data [23], the NPMI matrix reveals a prominent block-diagonal structure corresponding to known lineage-associated gene programs (Fig. 2A). An inset focusing on a submatrix representative of both lymphoid (T/NK) and myeloid marker genes shows strongly positive NPMI values within lineages, indicating consistent co-expression across cells (Fig. 2B). In contrast, cross-lineage gene pairs typically exhibit negative NPMI, reflecting transcriptional incompatibility rather than simple sparsity Fig. 2A, B). Gene pairs with NPMI values near zero show little enrichment or depletion of co-occurrence across cells, whereas NA values arise when no co-occurrence is observed in the dataset. These observations demonstrate that NPMI computed from binary presence–absence events recovers biologically meaningful co-expression patterns without requiring expression normalization or reference annotations.

**Figure 2.**
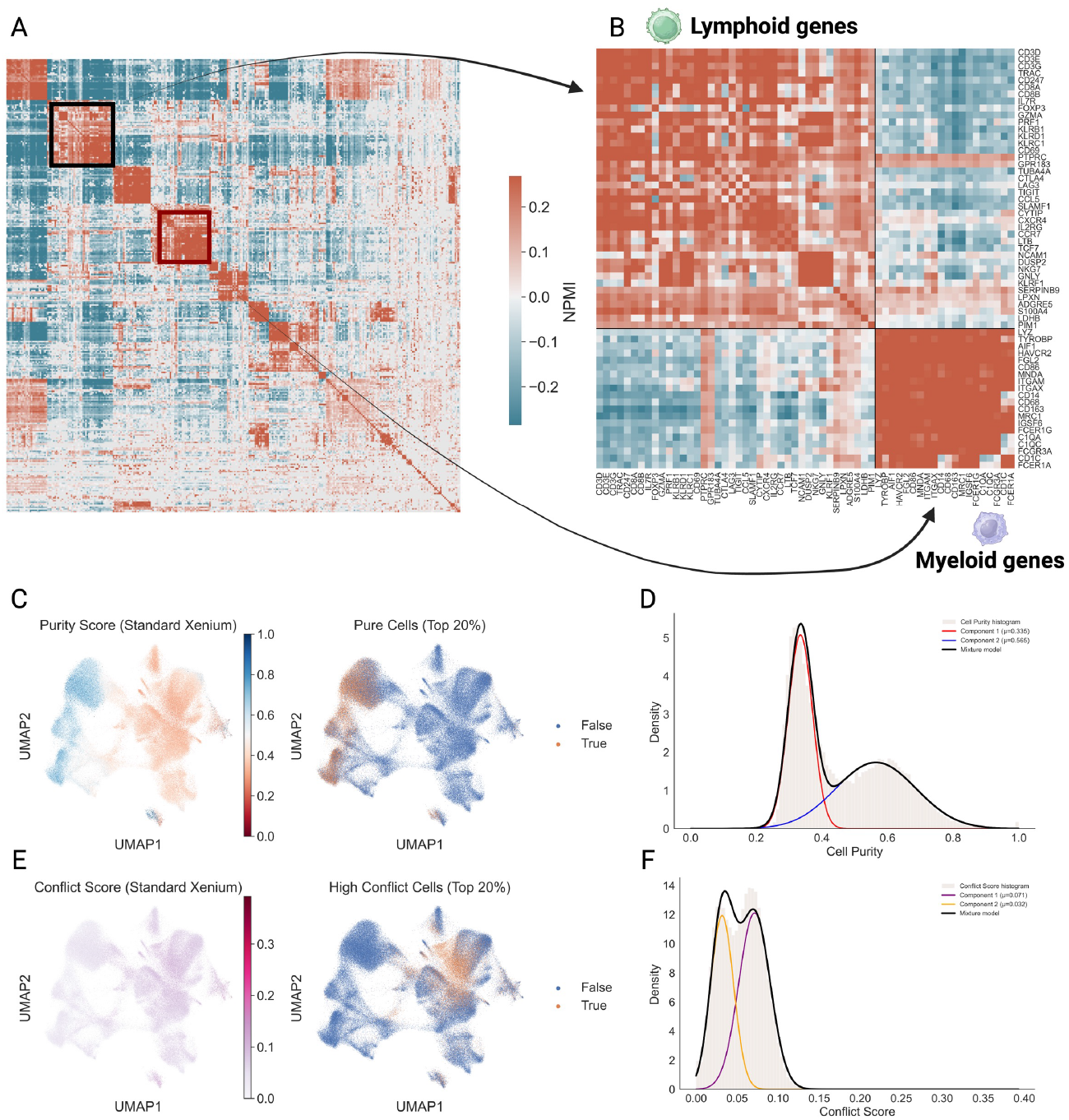
NPMI reveals lineage-resolved gene programs and enables quantitative purity and conflict scoring. (A) Global gene–gene NPMI matrix computed from the standard Xenium breast cancer dataset, showing block-diagonal lineage programs and negative cross-lineage associations. Insets highlight representative submatrices. (B) Representative lymphoid and myeloid gene modules with high within-lineage and negative cross-lineage NPMI. (D) UMAP embedding of whole-cell profiles colored by the NPMI-derived purity score, which quantifies internal gene–gene coherence. Cells in the top 20% of purity scores are indicated, marking transcriptionally coherent entities. (E) Distribution of purity scores across cells with a two-component Gaussian mixture model (GMM), revealing a bimodal structure separating high-purity from low-purity profiles. (F) UMAP embedding colored by the NPMI-derived conflict score, which captures mutually incompatible gene associations within individual cells. Cells in the top 20% of conflict scores localize predominantly to regions connecting distinct UMAP clusters. (G) Distribution of conflict scores and corresponding two-component GMM fit, separating low-conflict and high-conflict states associated with mixed-lineage transcriptional signatures.

Using this NPMI matrix, we next defined cell-level purity and conflict scores to quantify, respectively, the internal coherence and mutual incompatibility of gene programs within individual cellular profiles (see Methods). Purity and conflict scores, projected onto UMAP embeddings of Xenium breast cancer data, reveal distinct patterns Fig. 2C,E). Consistent with these patterns, both metrics exhibited bimodal distributions—confirmed by two-component Gaussian mixture models (GMM) [24]— separating coherent cells from ambiguous profiles Fig. 2D,F).

To assess the generality of these metrics across platforms and segmentation strategies, we applied the same purity and conflict scoring framework to Xenium lung cancer data with multimodal segmentation [25] and to Multiplexed Error-Robust Fluorescence in situ Hybridization (MERFISH) mouse ileum data segmented with Baysor using a 3D membrane prior [5]. In both datasets, cells with elevated conflict scores and reduced purity were again preferentially localized to narrow connective bands between UMAP clusters, rather than within well-defined cell-type clusters (Supplementary Fig. 2A,B; Supplementary Fig. 3A,B). These regions correspond to transcriptionally intermediate or mixed profiles, consistent with residual transcript sharing or boundary errors, as commonly observed for transitional cell states in low-dimensional embeddings of single-cell transcriptomic data [26–28].

Notably, in contrast to standard nucleus-expanded segmentation, we see fairly unimodal distributions of purity and conflict scores in the Xenium Lung Cancer and Baysor-segmented MERFISH data, as quantified by GMM (Supplementary Fig. 2C,D; Supplementary Fig. 3C,D). This unimodality relative to the standard Xenium segmentation could be attributed to the smaller area of the lung cancer tissue compared to the breast cancer tissue in the standard Xenium, and potential improvements in cell purity due to Baysor-segmentation. However, there are other confounding factors to consider such as the transcript selection and detection characteristics of each dataset. Together, these results indicate that NPMI-derived purity and conflict scores are reliable segmentation-agnostic indicators of transcriptional coherence and mixing across spatial transcriptomics technologies.

### 2.3 TRACER refines transcript assignment and restores cell-type resolution in the absence of multimodal staining

Based on the ability of the purity and conflict scores to detect transcriptional mixing, we next evaluated whether TRACER improves transcript assignment and downstream cell-type resolution (Fig. 3). We generated a version of the lung cancer dataset with nucleus-expanded cell segmentation. 2D convex hulls of the transcripts assigned to each cell are non-overlapping for both multimodal segmentation and nucleus-expanded segmentation(Fig. 3A,B), reflecting the strict boundary enforcement of the imaging-based approach. In contrast, TRACER-stitched and TRACER–fine-tuned outputs yield partially overlapping convex hulls (Fig. 3C,D), representing contributions from overlapping cells captured in the tissue section.

**Figure 3.**
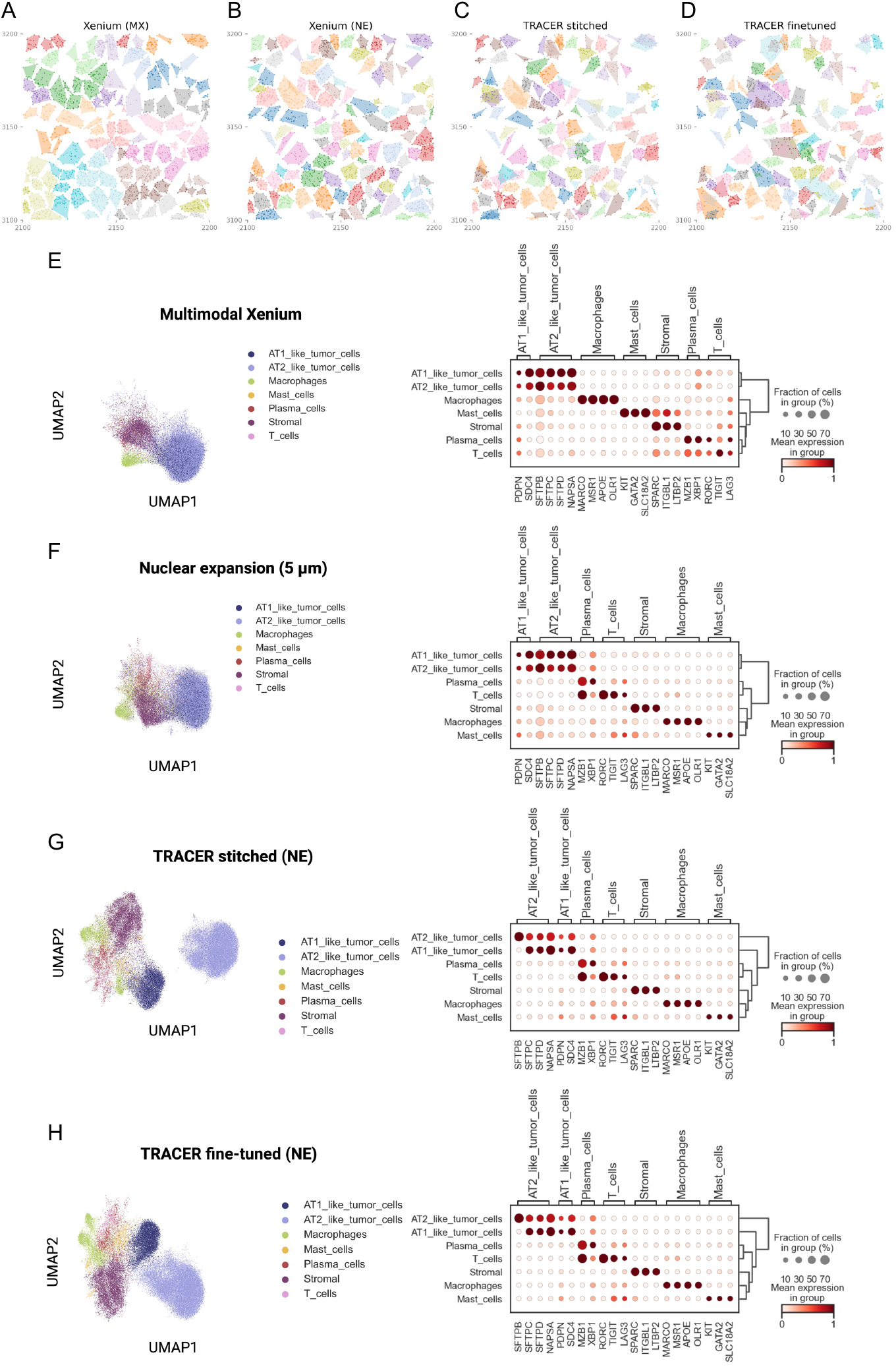
TRACER refines transcript assignment and cell-type resolution without multimodal staining. (A–D) Convex hulls generated from transcripts assigned to cells using (A) multimodal Xenium segmentation; (B) 5-*μ*m nuclear expansion (NE) without multimodal information, both showing strict boundary enforcement. (C) TRACER-stitched refinement applied to the NE baseline; (D) TRACER-fine-tuned refinement applied to the NE baseline, both allowing overlapping cells. Convex hulls are for illustrative purposes only. (E–H) UMAP embeddings and corresponding marker-gene dot plots for each segmentation strategy. (E) Multimodal Xenium segmentation; (F) 5-*μ*m NE without multimodal staining, showing cross-lineage spillover. (G) TRACER-stitched refinement applied to the NE baseline; (H) TRACER-fine-tuned refinement applied to the NE baseline, both exhibiting improved signal-to-noise ratio and clearer lineage separation.

UMAP embeddings based on multimodal segmentation or a 5 μm nuclear expansion showed substantial intermixing of lineages (Fig. 3E,F). Correspondingly, dot plots of canonical marker expression showed broad, low-specificity patterns, indicative of hybrid profiles due to transcriptional mixing. Applying TRACER to both segmentation outputs markedly improved manifold structure (Fig. 3G,H; Supplementary Fig. 2E,F): clusters become clearly delineated clusters, and marker expression profiles exhibit higher specificity with reduced cross-lineage contamination. Notably, TRACER applied to nuclear expansion-based segmentation achieved similar or improved cluster separation compared to multimodal segmentation.

TRACER produces deterministic outputs, yielding identical cellular reconstructions across runs (Supplementary Fig. 2G). At the individual cell level, TRACER increased relative purity and decreased relative conflict compared with pre-TRACER assignments (Supplementary Fig. 2H), consistent with reduced transcript mixing after refinement.

### 2.4 TRACER reveals partial cells arising from three-dimensional tissue geometry

Imaging-based spatial transcriptomics is intrinsically a 3D measurement problem. Most platforms acquire 3D transcript coordinates from tissue sections with non-trivial thickness[29–31]. As a result, 2D segmentation inevitably produces mixed profiles from boundary errors between overlapping adjacent cells. Partial cells with out-of-plane nuclei either contribute transcripts to overlapping in-plane cells—causing further contamination—or are missed entirely, their transcripts remaining unassigned (Fig. 1A).

TRACER addresses these artifacts by separating contaminated profiles and recovering missed partial cells. To formalize this, we define three categories of cells: whole, partial, and reconstructed partial cells. Whole cells are defined as cellular profiles containing a well-focused nucleus within the imaging plane, consistent with inclusion of the major cellular volume. Partial cells are fragments separated by TRACER from whole-cell profiles—representing contamination from overlapping cells with out-of-plane nuclei. Reconstructed partial cells are coherent transcript clusters assembled by TRACER from previously unassigned transcripts—representing potential cells that were missed entirely because they did not overlap with any detected nuclei (Fig. 1A).

To evaluate the biological relevance of these entities, we applied TRACER to the standard Xenium breast cancer dataset. UMAP embeddings from the original segmentation exhibited poorly separated clusters with extensive inter-cluster mixing (Supplementary Fig. 4A, left), consistent with transcript mixing and segmentation errors. TRACER-stitched and TRACER-fine-tuned whole-cell profiles formed compact, well-separated clusters (Supplementary Fig. 4A, middle and right). Projecting original study annotations onto these embeddings revealed that multiple major lineages—including DCIS tumor cells, invasive tumor cells, T cells, myeloid cells, myoepithelial populations, stromal cells, endothelial cells, and perivascular-like cells—were substantially conflated under the original segmentation (Supplementary Fig. 4B, left). After TRACER refinement, these lineages resolved into discrete clusters corresponding to biologically coherent cell types–using original annotations without re-annotating the cells (Supplementary Fig. 4B, middle and right).

To examine how TRACER resolves transcriptionally ambiguous profiles, we focused on a representative “T cell & tumor hybrid” cell from the original study (Fig. 4A)[23]. After TRACER refinement, transcripts from this single hybrid profile separate into two distinct, biologically coherent entities: one exhibiting a HER2-positive tumor gene program (Fig. 4B,C) and the other exhibiting a canonical CD8^+^ T cell program (Fig. 4D,E). We observed a similar effect for a cell from the cluster “unlabeled” in the original study (Supplementary Fig. 5A), where TRACER fine-tuning resolved a mixed profile into two coherent entities corresponding to a T cell–like program (Supplementary Fig. 5B,C) and a tumor-like program (Supplementary Fig. 5D,E).

**Figure 4.**
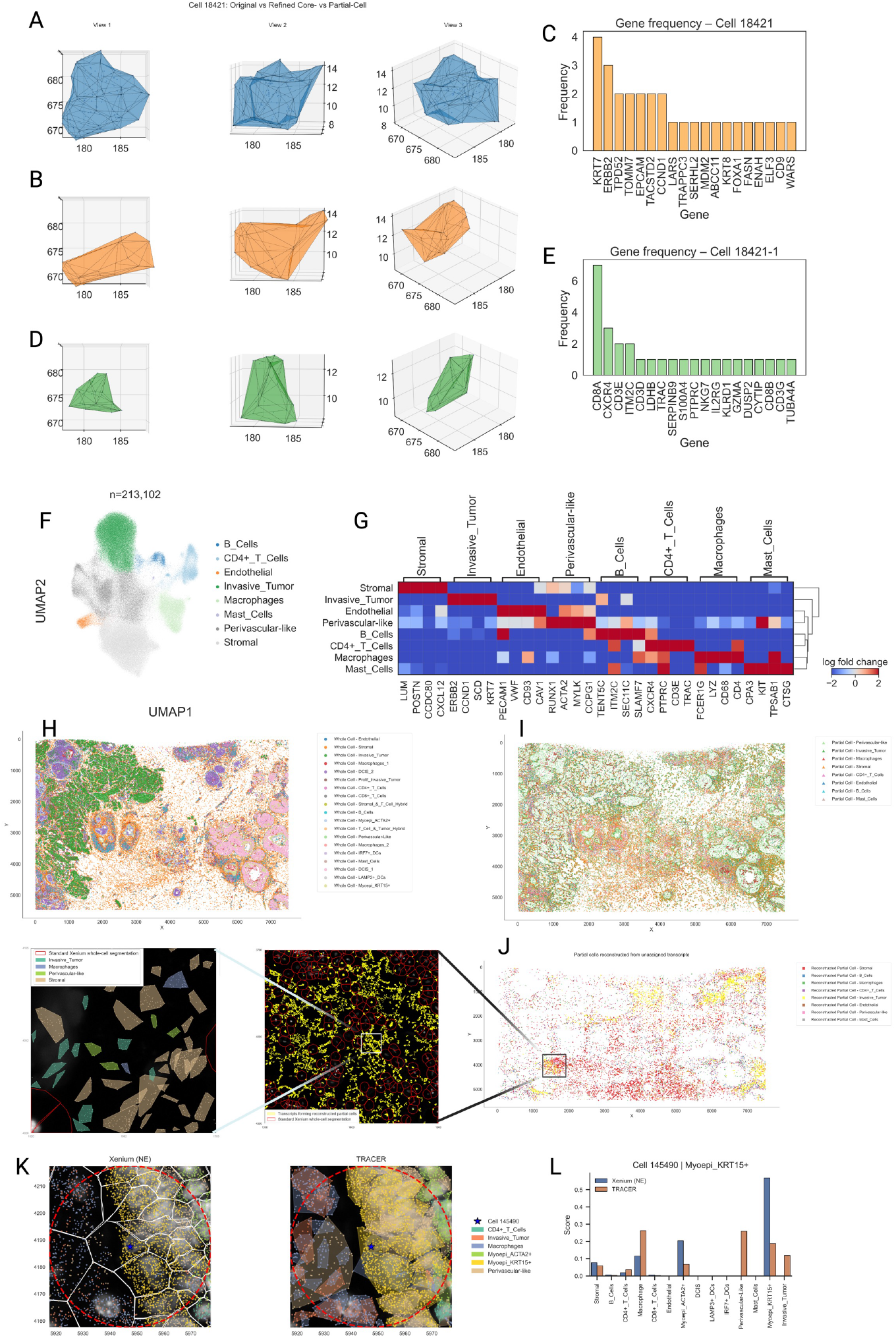
Recovery and characterization of partial cellular profiles with TRACER. (A) Three-dimensional concave hull constructed from the transcript point cloud originally assigned to a representative T cell–tumor hybrid cell under standard Xenium segmentation. (B,C) TRACER-extracted component corresponding to the tumor-associated subset of transcripts from the hybrid cell: (B) 3D concave hull and (C) transcript frequency profile. (D,E) TRACER-extracted component corresponding to the T cell–associated subset of transcripts: (D) 3D concave hull and (E) transcript frequency profile. (F) UMAP embedding of partial cells identified by TRACER after stitching and spatial fine-tuning. (G) Differential marker-gene expression matrix for Leiden clusters (annotated by marker-gene expression) computed from the partial cell population. (H) Spatial distribution of whole cells in the breast cancer Xenium section, colored by the original cell-type annotations. (I) Spatial distribution of TRACER-derived partial cells originating from segmentation-derived hybrid profiles, colored by Leiden clusters. (J) Spatial distribution of TRACER-derived partial cells reconstructed from previously unassigned transcripts, colored by Leiden clusters. The left inset shows transcripts contributing to reconstructed partial cells (yellow) overlaid on standard Xenium whole-cell segmentation boundaries (red). The right inset shows a convex-hull representation of these reconstructed partial cell transcripts within extracellular regions. (K,L) The inferred local neighborhood of a KRT15+ myoepithelial cell after including the TRACER-recovered partial cells in the vicinity shows increased cellular influence from macrophages, perivascular-like, and invasive tumor cells.

TRACER reconstructs partial cells by embedding unassigned transcripts in 3D physical space and representing them as nodes in a *k*-nearest-neighbor graph (Fig. 1A). The NPMI prior enforces internal transcriptional coherence within each reconstructed partial cell, allowing structured entities to emerge from the unassigned transcript pool rather than being discarded. UMAP embeddings of the partial and reconstructed partial cells reveal that these entities also form well-defined clusters. Unsupervised Leiden clustering of these partial cells [32] produced clear and biologically interpretable populations (Fig. 4F), each defined by distinct marker programs (Fig. 4G). For example, clusters expressed *LYZ* (macrophages), *LUM* (stromal cells), *CD3E* (T cells), *SLAMF7* (B cells), *ERBB2* (invasive tumor cells), *PECAM1* (endothelial cells), *ACTA2* (perivascular-like cells), or *KIT* (mast cells).

Spatially, the distribution of partial and reconstructed partial cells closely mirrors that of whole cells (Fig. 4H–J), consistent with vertical continuity of tissue architecture rather than artifactual reassignment. In Fig. 4J (inset), transcripts contributing to reconstructed partial cells were predominantly unassigned under standard segmentation and would have been discarded from downstream analysis. TRACER consolidates these transcripts into coherent reconstructed partial cells whose convex hulls fill the gaps between segmented cells (Fig. 4J, inset), further supporting their biological relevance. Inclusion of these partial cells in the cellular neighborhood analysis using FunCN [33] substantially changes the inferred cellular organization compared to a neighborhood that includes only whole cells, as demonstrated for a representative KRT15^+^ myoepithelial cell (Fig 4K,L).

### 2.5 TRACER resolves hybrid cellular profiles and removes segmentation-derived ligand–receptor artifacts

Because no gold-standard ground truth exists for evaluating segmentation accuracy in imaging-based spatial transcriptomics [15], we assessed TRACER performance indirectly through marker gene specificity—reflecting the primary purpose of segmentation, namely to capture biologically coherent cell types. In the Xenium breast cancer dataset, marker expression profiles show extensive cross-lineage contamination and weak cluster specificity (Supplementary Fig. 5F). In contrast, TRACER outputs yield markedly enhanced marker specificity and clearer lineage-appropriate expression patterns (Supplementary Fig. 5G; Fig. 5A), consistent with reduced transcript mixing and improved cell-type delineation.

**Figure 5.**
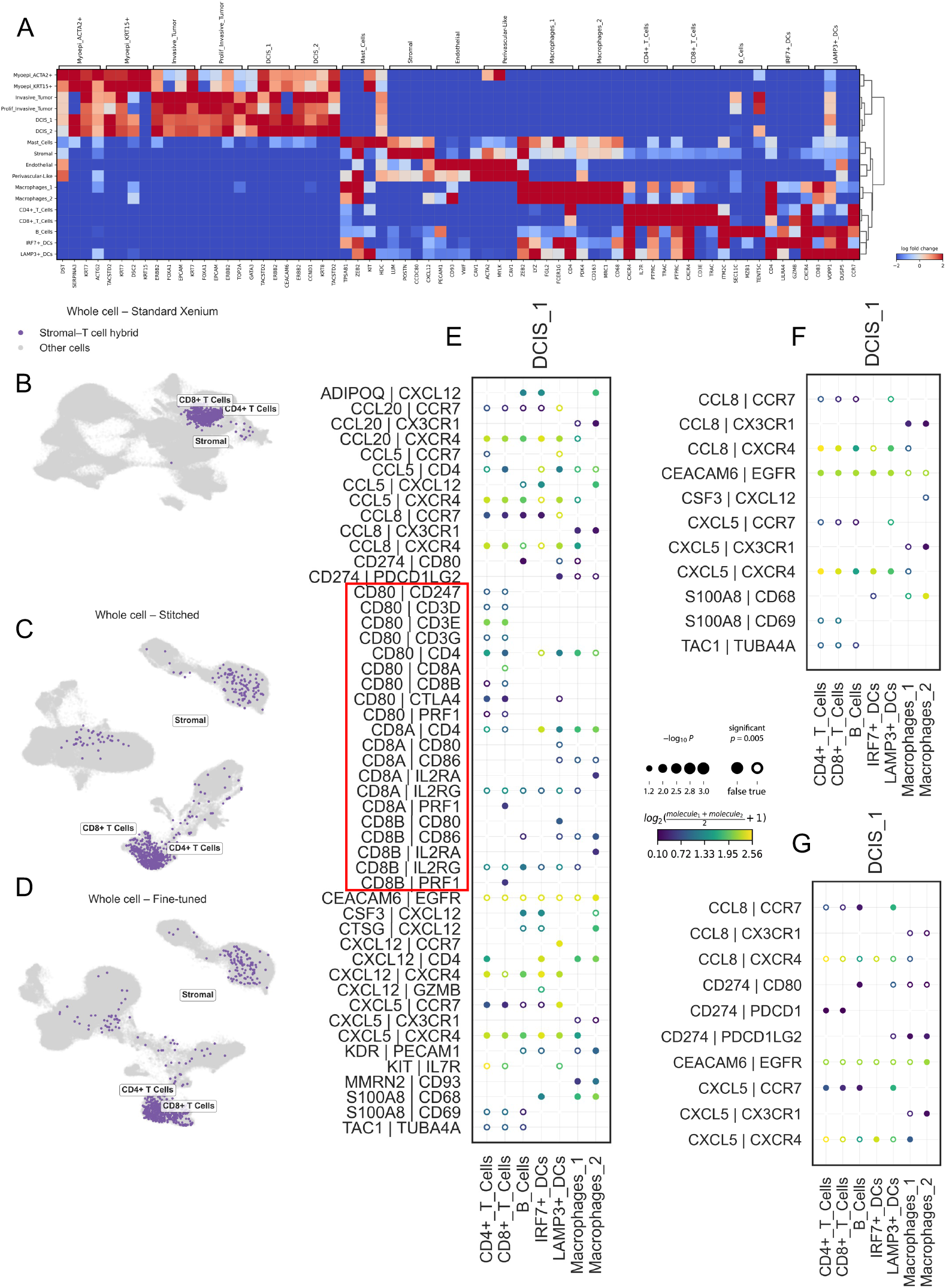
TRACER resolves hybrid cellular profiles and removes segmentation-derived ligand–receptor artifacts. (A) Differential marker-gene expression matrix for Leiden clusters derived from standard Xenium segmentation after TRACER fine-tuning. Sharpened lineage-specific profiles and reduced cross-lineage signal are observed. (B) UMAP embedding of cells derived from the standard Xenium segmentation, colored by original study annotations. A cluster annotated as “Stromal– T cell hybrid” forms a contiguous manifold and contains cells whose expression profiles include both stromal- and T-cell–associated markers. (C,D) UMAP embeddings of cells after TRACER refinement of the standard Xenium baseline. In TRACER-stitched (C) and TRACER–fine-tuned (D) embeddings, cells originally grouped within the hybrid manifold occupy two distinct regions corresponding to stromal-associated and T cell–associated expression profiles. No external annotations or marker-based constraints were used during TRACER processing; annotations are shown for visualization only. (E–G) Ligand–receptor permutation analysis (FDR *<* 0.05) between DCIS-1 and immune cell groups. Using standard Xenium segmentation (E), DCIS-1 cells appear to express immune-restricted ligands (e.g., CD80, CD8A, CD8B), producing unlikely interactions. These interactions are not observed when using TRACER-stitched (F) or TRACER-fine-tuned (G) transcript assignments, in which DCIS-1 retains only epithelial-associated ligand–receptor signals.

Next, we examined two prominent hybrid populations that arise from the segmentation in the original study. The first corresponds to a stromal & T cell hybrid cluster (Fig. 5B). After TRACER stitching and fine-tuning, transcripts from this hybrid population separate into distinct stromal and T cell clusters (Fig. 5C,D), restoring biologically expected boundaries. A similar pattern emerges in the T cell & tumor hybrid population identified in the original study, where transcriptional resolution separates them into distinct T cell and tumor clusters after TRACER refinement (Supplementary Fig. 4C). These findings illustrate TRACER’s ability to resolve mixed-lineage profiles produced by segmentation errors.

To illustrate the impact of segmentation errors on downstream analyses and TRACER’s ability to mitigate them, we compare the results of ligand-receptor interaction analysis using the outputs from the original segmentation and TRACER. The transcriptional mixing resulting from the original segmentation leads to the inference of biologically implausible ligand-receptor interactions [34, 35]. For example, in the original segmentation, ductal carcinoma in situ subtype 1 (DCIS-1) cells—previously characterized as low-invasive [23]—spuriously appeared to express immune-restricted ligands and receptors such as *CD80, CD8A*, and *CD8B*, result in unlikely DCIS-to-immune cell interactions (Fig. 5E). TRACER refinement substantially reduces these likely artifacts, yielding interaction patterns more consistent with lineage-restricted ligands and receptors and clarifying epithelial–immune communication patterns. These corrections illustrate the downstream effect of TRACER for suppressing spurious ligand–receptor interaction predictions arising from transcript mixing from segmentation-errors.

### 2.6 Quantitative benchmarking of segmentation fidelity using coherence-based metrics

We benchmarked TRACER against multiple segmentation approaches across platforms and datasets using a combination of coherence-based and complementary quality metrics (Fig. 2; Supplementary Figs. 2–3). Coherence-based metrics included NPMI-derived purity and conflict scores, which quantify the extent of coherent and incompatible intracellular gene–gene associations, as well as relative purity and relative conflict, defined as the fractions of total intracellular association signal attributable to coherent and conflicting gene pairs, respectively. As complementary measures, we defined signal strength as the total magnitude of positive and negative NPMI associations within each cell and retained transcript count per cell as an orthogonal quality-control metric.

We first evaluated TRACER on the MERFISH mouse ileum dataset introduced in the Baysor study, in which volumetric segmentation was guided by Cellpose-derived 3D membrane priors [5]. Compared with the original Baysor segmentation, TRACER-stitched whole cells showed a marked increase in purity, with a further improvement after fine-tuning, together with reduced conflict (Fig. 6A; Supplementary Fig. 3I). Transcript counts decreased modestly after TRACER refinement, whereas signal strength decreased more substantially, indicating that the transcripts removed by TRACER were disproportionately associated with incoherent gene–gene relationships rather than dominant expression programs. By contrast, Proseg [36] produced cellular profiles with transcript counts closer to those observed in TRACER-derived partial cells. Notably, although Baysor yielded higher transcript counts than Cellpose, consistent with the original report [5], these profiles exhibited lower purity and higher conflict in our coherence-based analysis (Fig. 6A; Supplementary Fig. 3I). Similar trends were observed in the standard Xenium breast cancer dataset and in both multimodal and nucleus-expanded Xenium lung cancer datasets, where TRACER-derived whole cells consistently showed higher purity and lower conflict than Baysor, Proseg, BIDCell [37] and Segger [38] (Supplementary Fig. 4D; Supplementary Fig. 7A).

**Figure 6.**
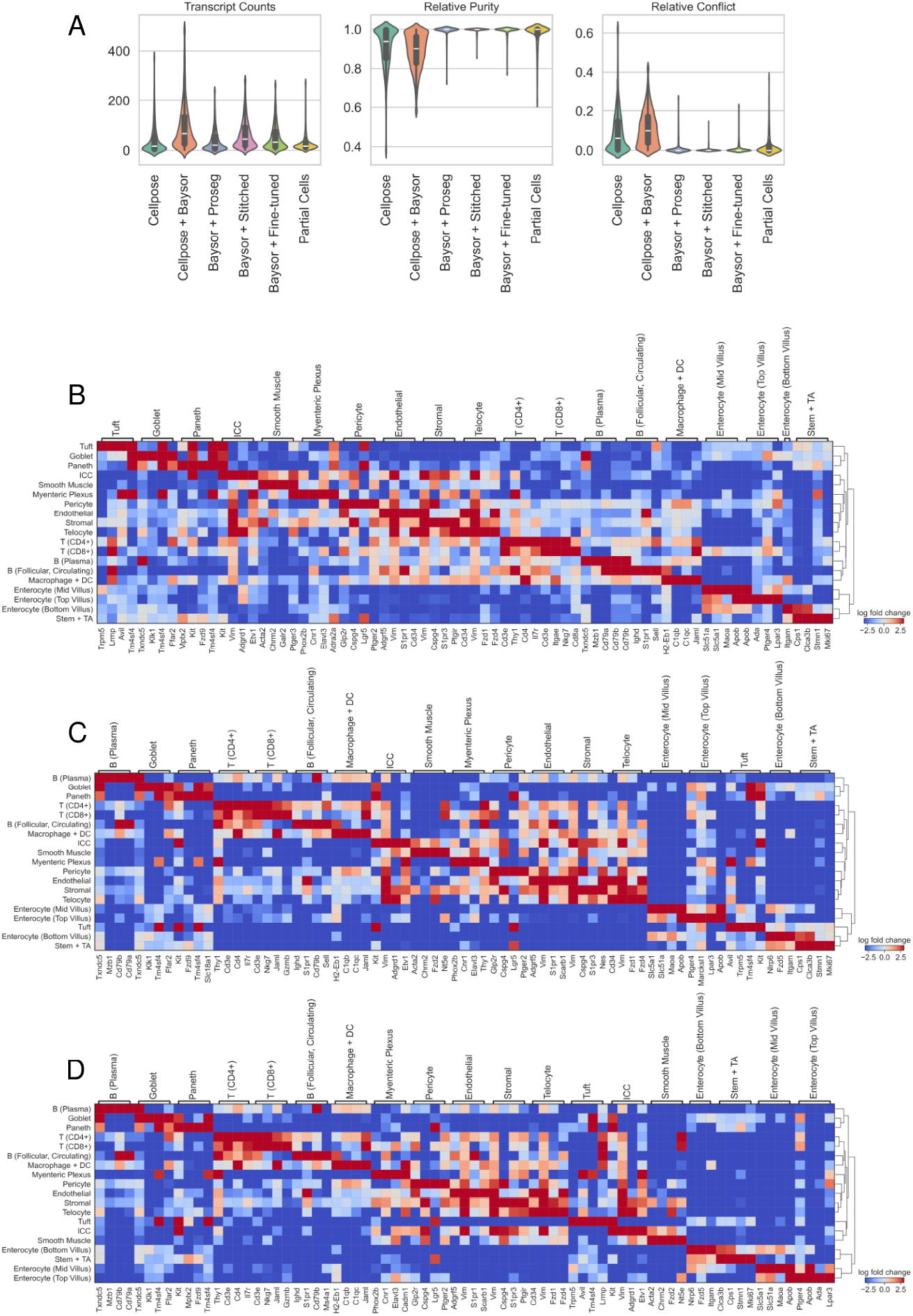
Quantitative benchmarking of TRACER relative to 3D membrane– guided segmentation. (A) Distributions of transcript counts, NPMI-derived relative purity and relative conflict for cellular entities obtained with Cellpose, Baysor using Cellpose-derived 3D membrane priors, Proseg using Baysor as a 3D prior, TRACER-stitched using Baysor as input, TRACER-fine-tuned using Baysor as input, and TRACER-derived partial cells. Relative purity and relative conflict quantify the fractions of positive and negative gene–gene associations within each cellular profile, respectively. TRACER-derived whole cells show higher relative purity and lower relative conflict than Baysor-segmented cells. (B) Differential marker-gene expression matrix for cell types annotated in the Baysor-segmented mouse ileum dataset, representing the transcriptional organization obtained from a 3D membrane–guided reference segmentation. (C,D) Differential marker-gene expression matrices for cell groups identified using TRACER-stitched (C) and TRACER-fine-tuned (D) outputs. Despite no cell-type annotations or marker genes were used during TRACER refinement, TRACER output exhibit improved signal-to-noise ratio relative to Baysor. Annotations are shown for visualization only.

We also compared the fidelity of the whole-cell profiles generated by Baysor and TRACER (Supplementary Fig. 6) using Scrublet, a widely used scRNA-seq doublet detector [39]. Although Scrublet was not developed for imaging-based spatial transcriptomics, segmentation-induced transcript mixing manifests as “doublet-like” profiles in expression space [40]. Baysor-segmented cells exhibited elevated Scrublet scores concentrated in the cluster-bridge regions highlighted by NPMI-derived conflict metrics, consistent with mixed-lineage transcript contamination. Across all expected doublet-rate settings, TRACER-stitched and TRACER-fine-tuned segmentations showed near-zero predicted doublets and uniformly low Scrublet scores, indicating that TRACER restores gene–gene coherence and eliminates mixed-lineage artifacts.

Beyond metric-level comparisons, TRACER also improved downstream structure and interpretability. In the MERFISH ileum dataset, unsupervised Leiden clustering of TRACER-corrected profiles produced more clearly separated UMAP clusters than those obtained from the original Baysor segmentation (Supplementary Fig. 3E), while remaining consistent with the annotated cell types from the original study (Supplementary Fig. 3F). Differential marker gene expression analysis further revealed enhanced signal-to-noise ratios following TRACER stitching and fine-tuning (Fig. 6B–D). In addition, TRACER identified 3,275 partial cells in this dataset (Supplementary Fig. 3G), which formed distinct clusters with coherent marker gene-expression programs (Supplementary Fig. 3H), reinforcing the biological interpretability of TRACER-derived entities.

## 3 Discussion

Accurate cell segmentation is a foundational requirement for faithful representation of cell types and cell states in imaging-based spatial transcriptomics. However, these segmentation errors arise from applying 2D methods to inherently 3D tissues, leading to hybrid clusters and lineage-mixed profiles. These errors lead to transcript mixing, substantially distorting downstream analyses including cell clustering, marker gene interpretation, cell-type identification, and ligand–receptor interaction analysis.

Recent computational approaches increasingly apply graph neural networks (GNNs) or contrastive learning frameworks to transcript–transcript or cell–cell graphs [9, 38]. While powerful, these approaches typically involve model training and greater computational requirements, with less transparent decision rules. In contrast, TRACER adopts a transparent, deterministic formulation: it computes NPMI representing gene-gene association directly from nucleus-derived contexts in the data and uses these values to guide coherence-based pruning and hierarchical stitching. TRACER requires no neural architectures and thus no model learning. This deterministic structure ensures reproducibility and allows users to inspect and validate every stage of the refinement process. Supplementary Table 1 summarizes the runtime and memory characteristics of TRACER across three benchmark datasets, demonstrating that the full pipeline can be executed efficiently on standard hardware without GPU acceleration.

In addition to TRACER, this work introduces NPMI-derived coherence metrics at the cellular level—purity, conflict, and related quantities—as segmentation-agnostic diagnostics for evaluating transcriptional mixing. These address a critical gap: imaging based segmentation benchmarks cannot capture the fuzzy, overlapping, and functionally defined nature of cell boundaries [15]. Coherence-based metrics provide a platform-independent approach to evaluate assess the biological coherence of cellular profiles, rather than geometric segmentation accuracy. We anticipate that these metrics will be broadly useful for algorithm development, benchmarking, and routine quality control in spatial transcriptomics.

TRACER also reframes partial cells as natural consequences of sectioning a 3D tissue structure, rather than as noise to be discarded. Through modeling, simulation, and empirical analysis, we show that these entities encode biologically meaningful gene programs, spatial organization, and tissue-level structure when treated explicitly. Recent work has independently demonstrated the biological relevance of these unassigned transcript populations [41]. Supplementary Table 2 summarizes how transcripts are reassigned across TRACER stages (pruning, stitching, fine-tuning) into whole cells, partial cells, and reconstructed partial cells across all benchmark datasets. While these entities represent incomplete cellular profiles and should not be used to define fine-grained cell states, they offer valuable information about tissue architecture and spatial niches.

At the same time, important limitations remain. Like many computational methods, TRACER requires parameter tuning, such as the maximum spatial distance used during fine-tuning. Increasing this threshold can merge spatially distant transcripts into overly large partial cells (Supplementary Fig. 7B–E). Although our recommended default thresholds performed consistently across tissues and platforms, future work integrating adaptive or data-driven distance estimation may further improve robustness. Computing NPMI from noisy input segmentation may yield misleading gene associations, and undersampling of rare genes may create spurious mutual exclusion—an inherent issue with data-driven methods. Furthermore, population-level statistics may not capture rare transitional states, such as cells undergoing epithelial–mesenchymal transition or macrophage repolarization, which may inadvertently be pruned by TRACER despite legitimately expressing markers that are negatively associated at the population level. To mitigate these issues, we computed NPMI from nuclear segmentations, which are less susceptible to boundary errors than whole-cell segmentations, and observed no obvious artifacts. We also adopted a conservative pruning strategy: using a moderate NPMI threshold and ignoring evidence of perfect mutual exclusion between rare gene pairs. Future work could address these limitations by incorporating external data sources such as single-cell RNA-seq atlases for NPMI computation, confidence intervals to account for gene frequency and estimation uncertainty, and trajectory-informed coherence metrics to preserve transitional populations.

Nevertheless, the modular design of TRACER enables independent modification of each stage—co-expression estimation, pruning, stitching, hierarchical assembly, and spatial fine-tuning—without restructuring the overall pipeline. As imaging-based spatial transcriptomics moves toward thicker tissues and true 3D acquisition, coherence-based refinement of cellular profiles will become increasingly critical.

## 5 Supplementary Materials

**Supplementary Figure 1.**
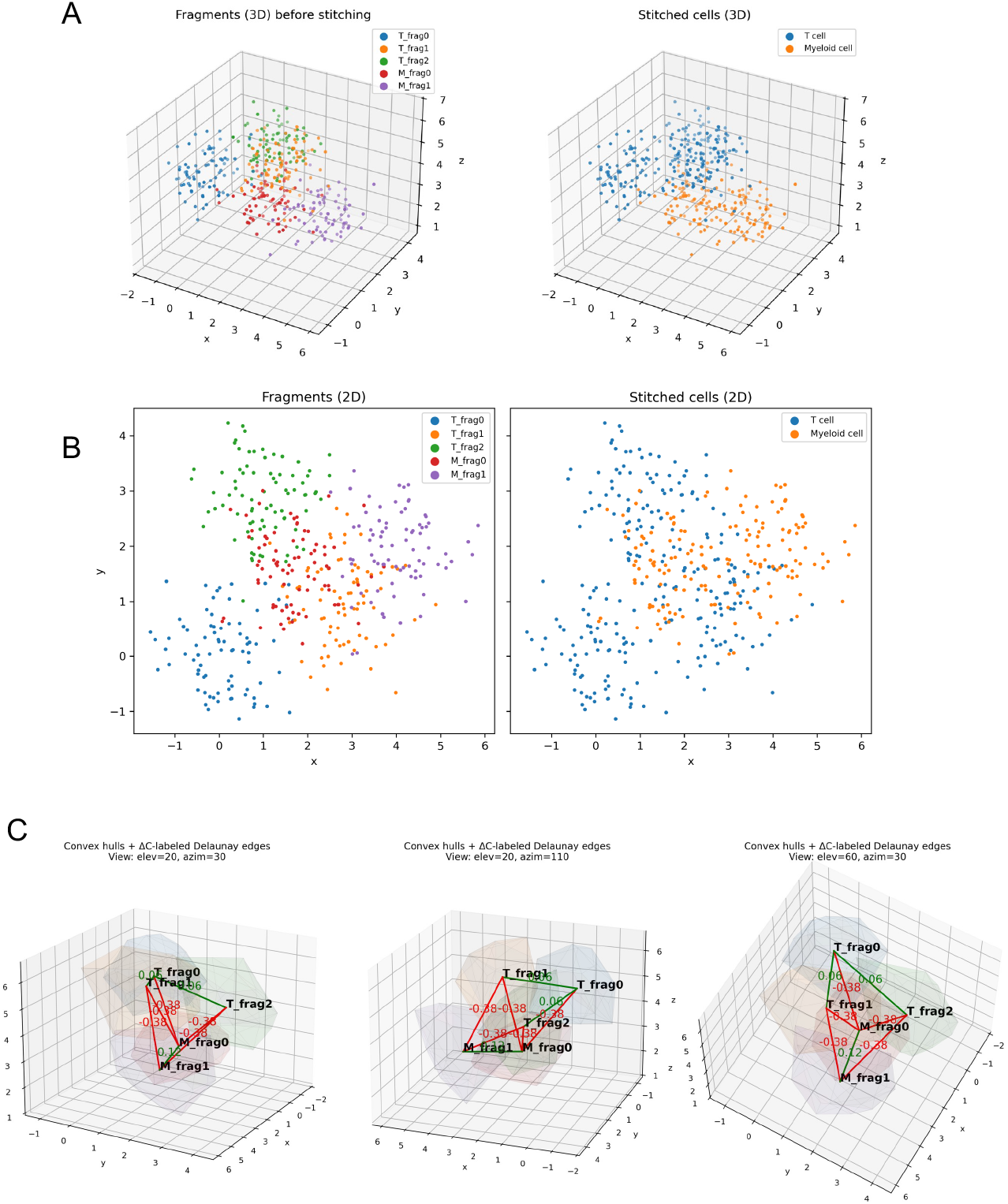
Δ*C*-guided stitching resolves fragmented cellular profiles in a simulated 3D system. (A) Simulated three-dimensional transcript point clouds representing five fragments (T_frag0, T_frag1, T_frag2, M_frag0, M_frag1) derived from two underlying cells (one T cell and one myeloid cell). (B) Two-dimensional projections of the same fragments, showing substantial spatial overlap that would confound merging strategies based solely on 2D segmentation strategies. (C) Δ*C*-based coherence-guided stitching applied to all fragment pairs. Candidate merges are evaluated by the change in transcriptional coherence (Δ*C*), enabling selective merging of fragments originating from the same cell while rejecting merges between transcriptionally incompatible fragments. Edge colors denote Δ*C* values for each candidate pair, illustrating coherent (positive Δ*C*) versus incoherent (negative Δ*C*) associations.

**Supplementary Figure 2.**
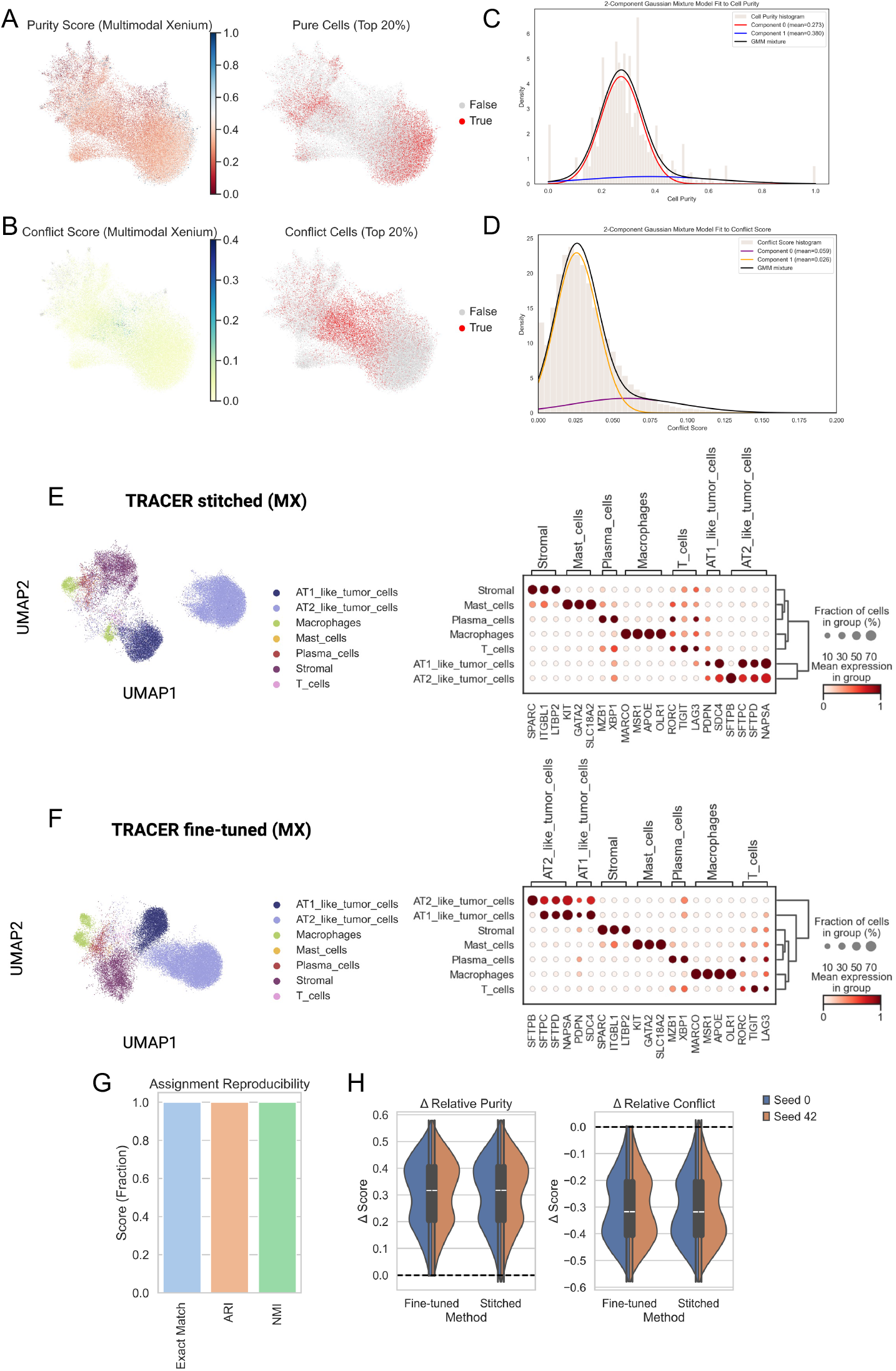
TRACER refinement and purity/conflict profiling in multimodal Xenium lung cancer data. (A) UMAP embedding of whole-cell profiles from a multimodal Xenium (MX) lung cancer dataset colored by the NPMI-derived purity score. Cells within the top 20th percentile of the purity distribution are highlighted. (B) UMAP embedding colored by the NPMI-derived conflict score, with the top 20th percentile of high-conflict cells highlighted. (C,D) Distributions of purity (C) and conflict (D) scores fitted using two-component Gaussian mixture models (GMMs). (E,F) UMAP embeddings and corresponding marker-gene dot plots for TRACER refinements applied to the multimodal Xenium segmentation. (E) TRACER-stitched refinement. (F) TRACER-fine-tuned refinement. (G) Assignment reproducibility across two independent TRACER runs with different random seeds (0 and 42), quantified using exact-match fraction, adjusted Rand index (ARI), and normalized mutual information (NMI). TRACER assignments are identical across runs (exact-match fraction = 1.0). (H) Per-cell changes in relative purity and relative conflict across the two runs, with refinement-associated shifts toward higher purity and lower conflict at the per-cell level.

**Supplementary Figure 3.**
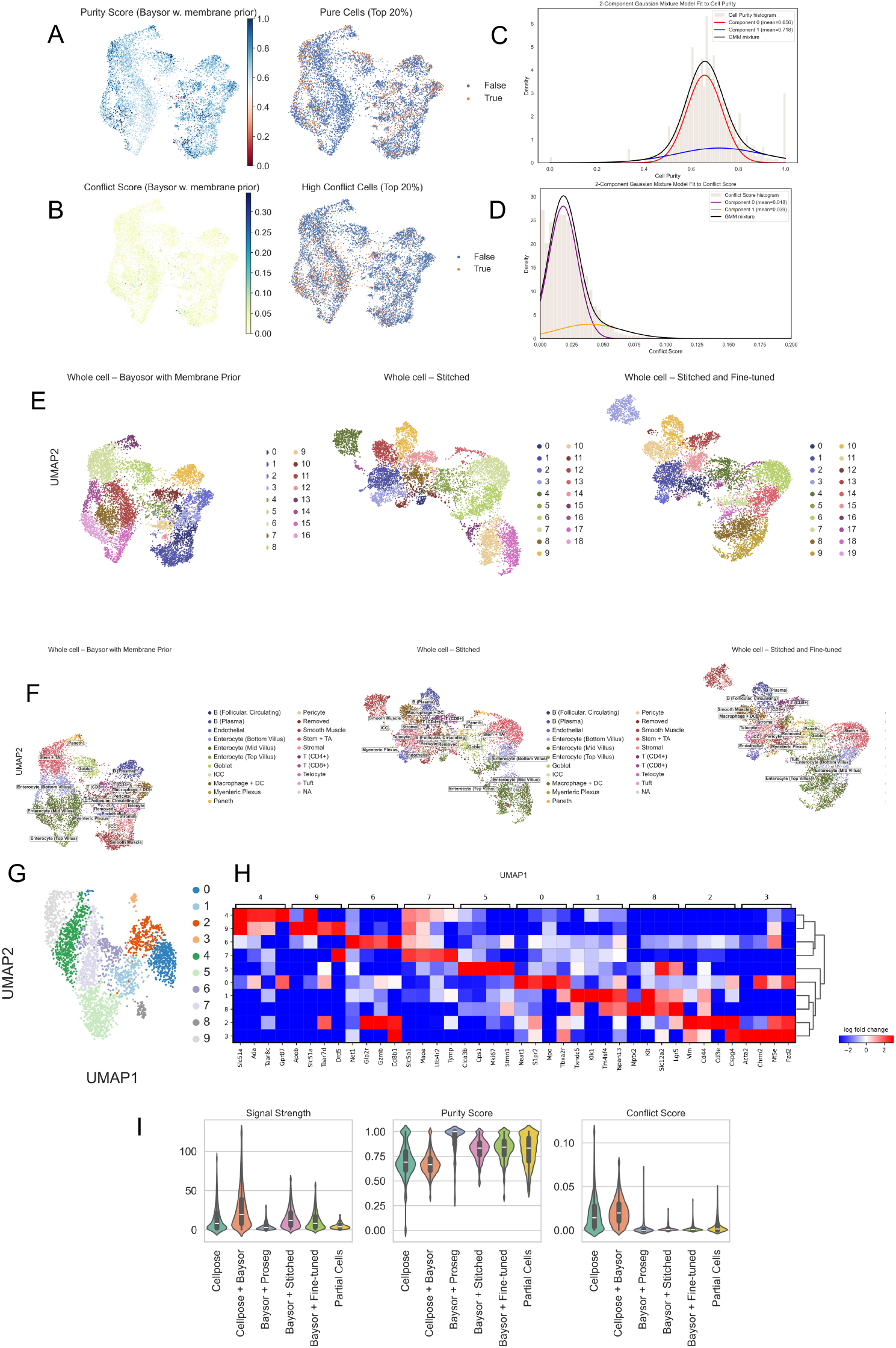
Purity/conflict scoring, TRACER refinement, and coherence metrics for MERFISH mouse ileum segmentation with membrane-informed 3D priors. (A) UMAP embedding of whole-cell profiles obtained from Baysor segmentation incorporating Cellpose-derived three-dimensional membrane priors, colored by NPMI-derived purity scores. Cells in the top 20th percentile of purity are highlighted. (B) UMAP embedding colored by NPMI-derived conflict scores, with high-conflict cells (top 20th percentile) highlighted. (C,D) Distributions of purity (C) and conflict (D) scores fitted with two-component Gaussian mixture models (GMMs). (E) Leiden clustering overlaid on UMAP embeddings for Baysor + membrane priors (left), TRACER-stitched whole cells (middle), and TRACER–fine-tuned whole cells (right). (F) Cell-type annotations from the original study overlaid on the corresponding embeddings in (E). (G) UMAP embedding of TRACER-identified partial cells after stitching and spatial fine-tuning, colored by Leiden cluster identity. (H) Differential marker-gene expression matrix for Leiden clusters shown in (G). (I) Distributions of signal strength and NPMI-derived purity and conflict across cellular entities generated by Baysor with Cellpose-derived membrane priors, Proseg with Baysor-derived priors, TRACER-stitched, TRACER–fine-tuned and TRACER-derived partial cells. TRACER-derived whole cells show increased purity and reduced conflict relative to Baysor-segmented cells.

**Supplementary Figure 4.**
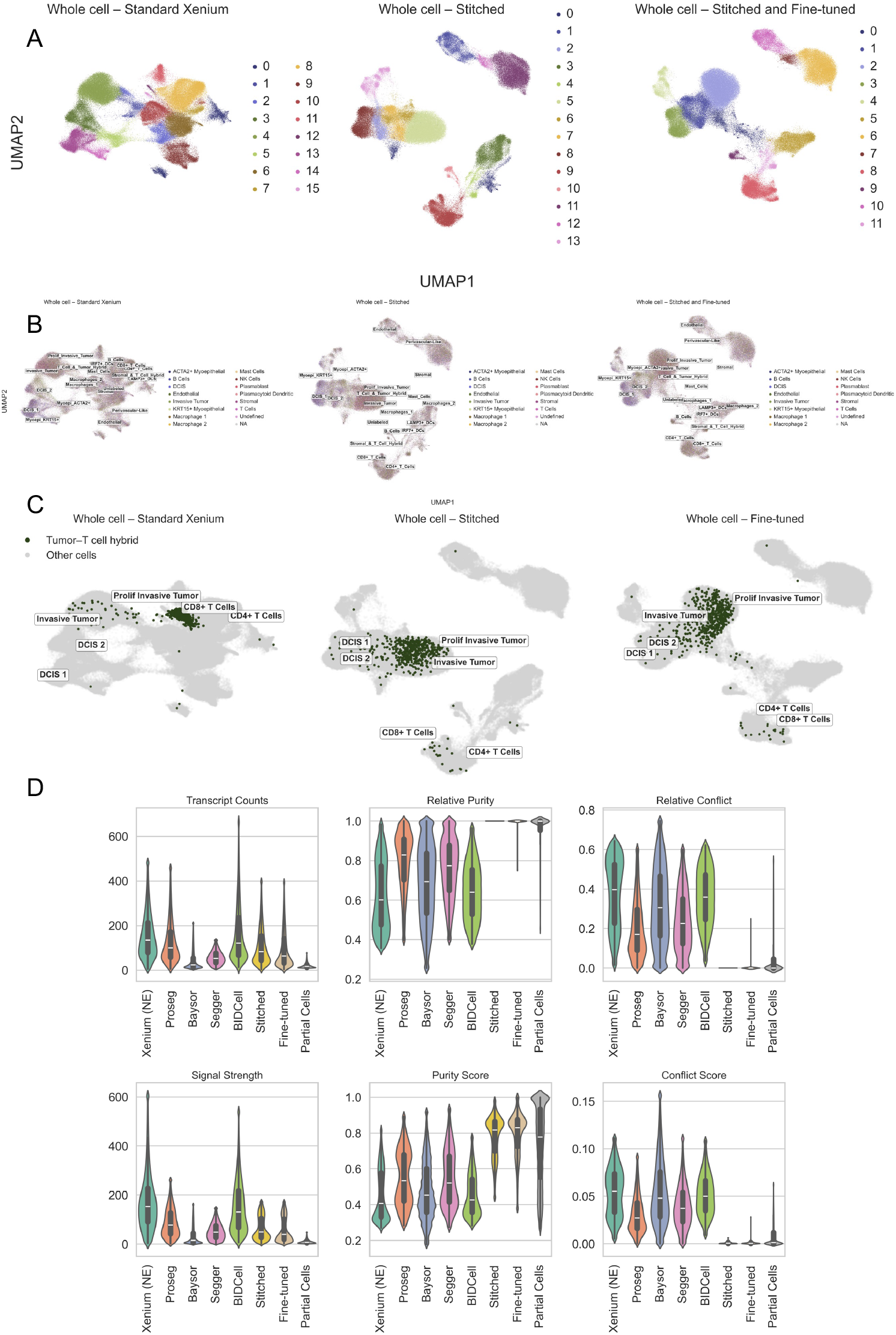
TRACER-induced changes in clustering structure, cell-type separation, and coherence metrics in standard Xenium breast cancer data. (A) Leiden clustering overlaid on UMAP embeddings for the standard Xenium segmentation (left), TRACER-stitched whole cells (middle), and TRACER– fine-tuned whole cells (right). (B) Cell-type annotations from the original study overlaid on the corresponding embeddings in (A). These labels were not used in TRACER and are shown for visualization only. (C) Distribution of cells originally annotated as a tumor–T cell hybrid population. Under the original segmentation (left), these cells occupy a shared manifold. After TRACER stitching (middle) and subsequent fine-tuning (right), the same cells separate into distinct tumor and T cell manifolds. (D) Distributions of transcript counts, signal strength, relative purity, relative conflict, and NPMI-derived purity and conflict across cellular entities from Xenium nucleus-expanded segmentation (Xenium (NE)), Proseg, Baysor, Segger, BIDCell, TRACER-stitched, TRACER–fine-tuned and TRACER-derived partial cells. Relative purity and relative conflict denote the fractions of coherent and incompatible gene–gene associations within each profile, respectively, and signal strength reflects the total magnitude of intracellular NPMI associations. TRACER refinement increases purity and reduces conflict across input segmentations.

**Supplementary Figure 5.**
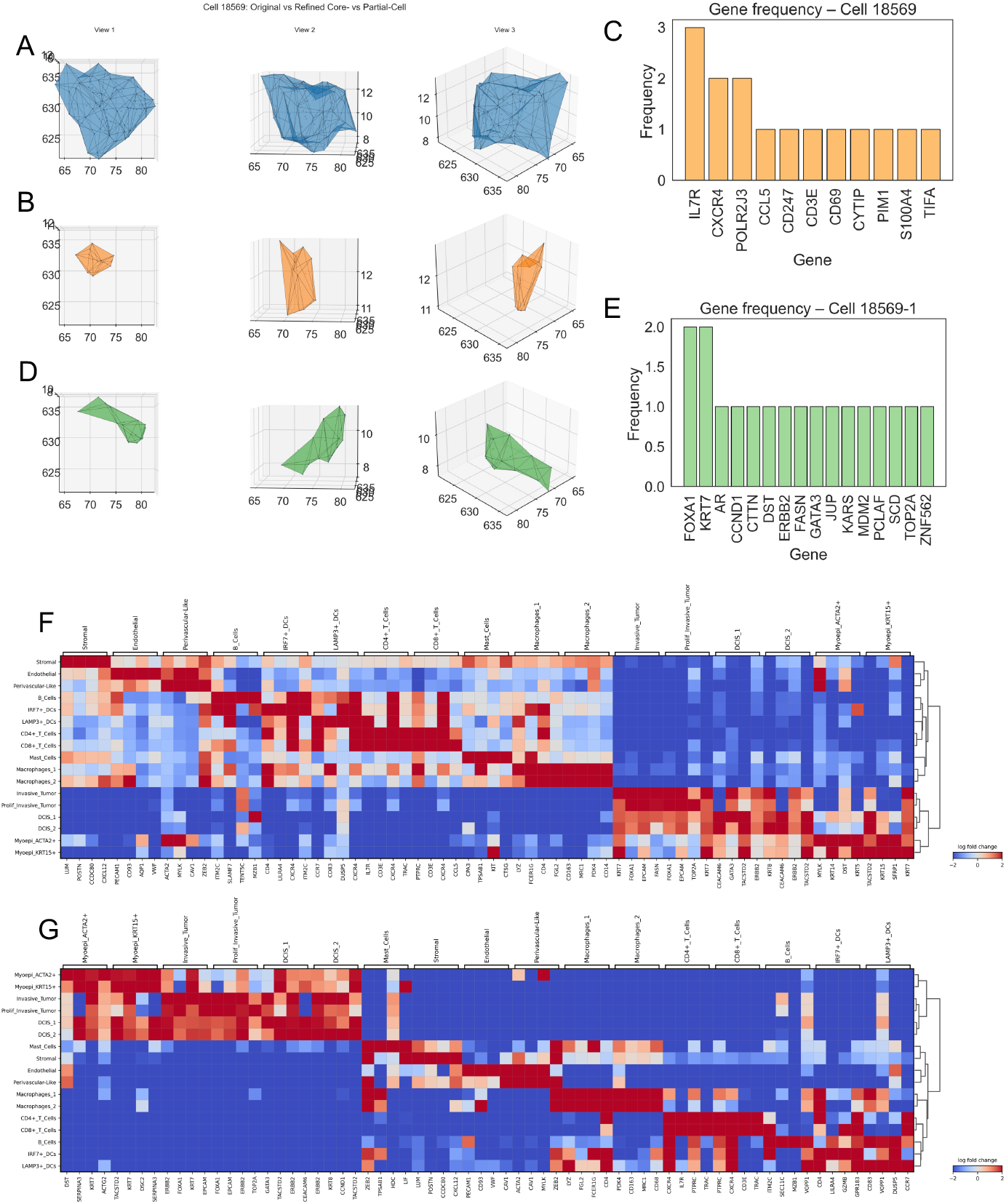
Resolution of mixed-lineage cellular profiles and coherence-guided decomposition of ambiguous cells. (A) Three-dimensional concave hull constructed from the transcript point cloud originally assigned to an unlabeled cell under the standard Xenium segmentation. This cell displayed promiscuous marker gene expression, preventing confident assignment in the original study. Three orthogonal views of the reconstructed concave hull are shown. (B,C) TRACER-derived component corresponding to a T cell, represented as a 3D concave hull (B) together with its gene frequency profile (C). (D,E) TRACER-derived component corresponding to a tumor cell, represented as a 3D concave hull (D) and its gene frequency profile (E). (F) Differential marker gene expression matrix for cell-type annotations defined by the original study applied to the standard Xenium segmentation, showing mixed-lineage signal across annotated groups. (G) Differential marker gene expression matrix for the same published cell-type annotations applied to TRACER-stitched correction, showing more distinct lineage-associated marker expression. Cell-type labels are derived entirely from the original peer-reviewed dataset and were not used during TRACER refinement.

**Supplementary Figure 6.**
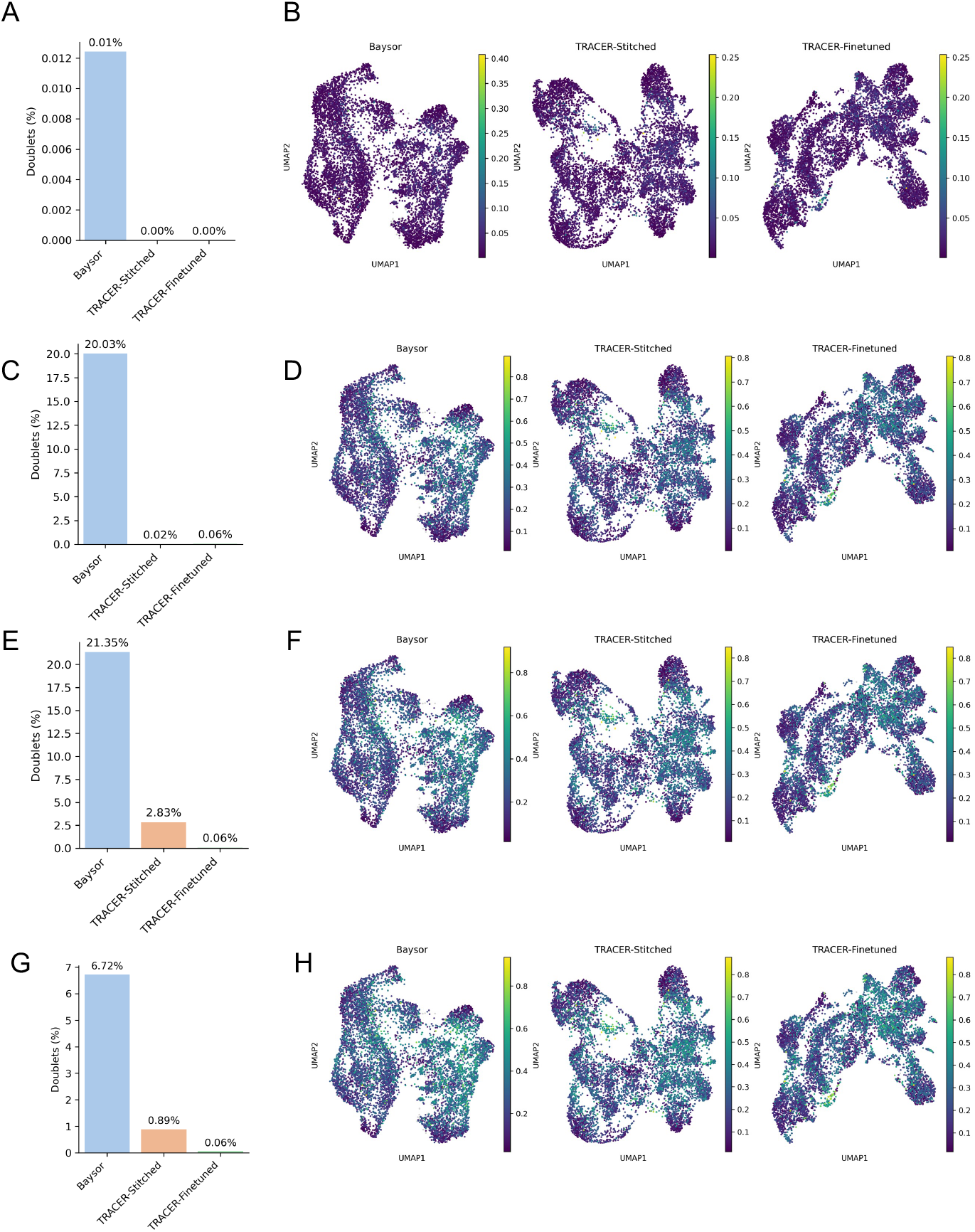
Scrublet-based expression doublet scores across segmentation strategies. (A–B) Scrublet applied with an expected doublet rate of 0.02. (A) Predicted doublet percentages for Baysor, TRACER-stitched, and TRACER-fine-tuned whole-cell profiles. (B) UMAP embeddings colored by Scrublet doublet scores. (C–D) Scrublet predictions using an expected doublet rate of 0.20. (E–F) Scrublet predictions using an expected doublet rate of 0.25. (G–H) Scrublet predictions using an expected doublet rate of 0.30. Across expected doublet priors, TRACER-derived cellular profiles exhibit consistently near-zero predicted doublet fractions and remain stable across Scrublet settings, whereas Baysor-derived profiles show substantially higher predicted doublet rates and stronger sensitivity to the assumed doublet prior.

**Supplementary Figure 7.**
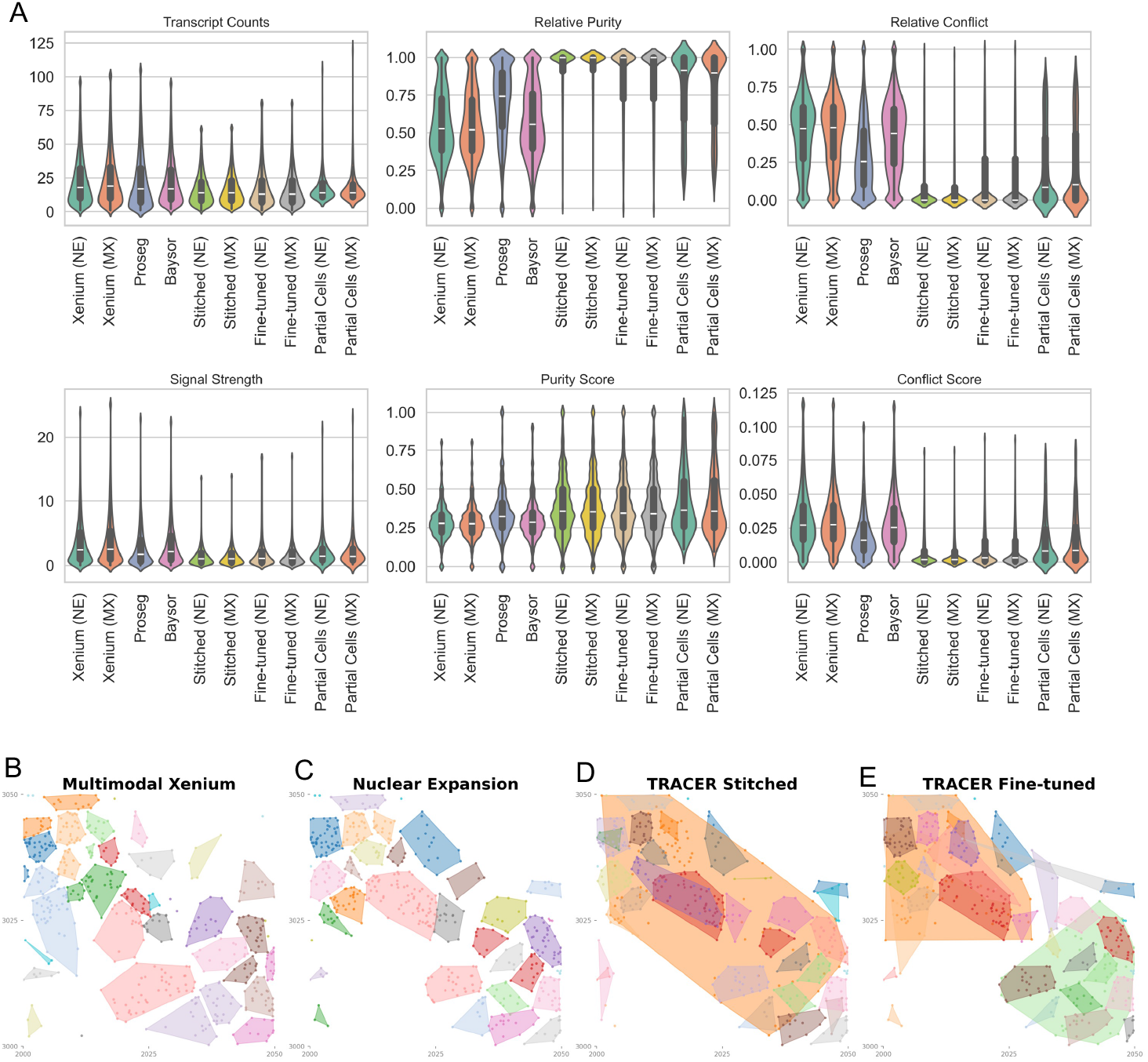
Summary coherence metrics and convex-hull visualizations for lung cancer multimodal Xenium across segmentation strategies. (A) Distributions of transcript counts, signal strength, relative purity, relative conflict, and NPMI-derived purity and conflict across cellular entities generated by Xenium nucleus-expanded (5-*μ*m) segmentation (Xenium (NE)), multimodal Xenium segmentation (Xenium (MX)), Proseg, Baysor, TRACER-stitched whole cells, TRACER-fine-tuned whole cells, and TRACER-derived partial cells from both NE and MX inputs. Relative purity and relative conflict denote the fractions of coherent and incompatible gene–gene associations within each profile, respectively, and signal strength reflects the total magnitude of intracellular NPMI associations. TRACER-derived whole cells show increased purity and reduced conflict relative to the input segmentations. (B–E) Comparison of convex hulls constructed from transcript-to-cell assignments across segmentation methods. (B) Multimodal Xenium segmentation, (C) 5-m nuclear-expansion segmentation lacking multimodal information, (D) TRACER-stitching of the nuclear-expansion baseline with relaxed spatial constraints (15 m), and (E) TRACER fine-tuned refinement of the nuclear-expansion baseline with relaxed spatial constraints (15 m). Note: relaxed spatial constraints in (D) and (E) can yield implausibly large convex hulls.

**Supplementary Table 1.**
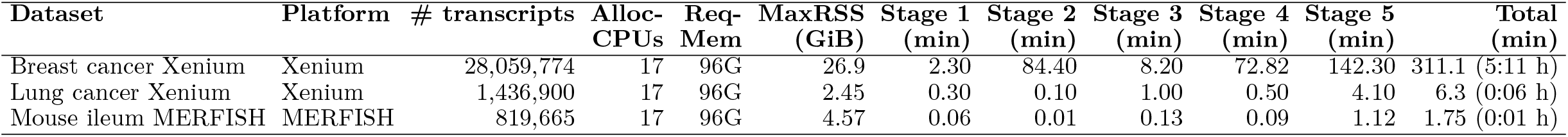
TRACER runtime and memory usage on three benchmark datasets. Wall-clock time was measured via SLURM (`sacct`); per-stage runtimes were obtained from internal logs. All jobs used 17 allocated CPU cores and 96 GB requested memory.

**Supplementary Table 2.**
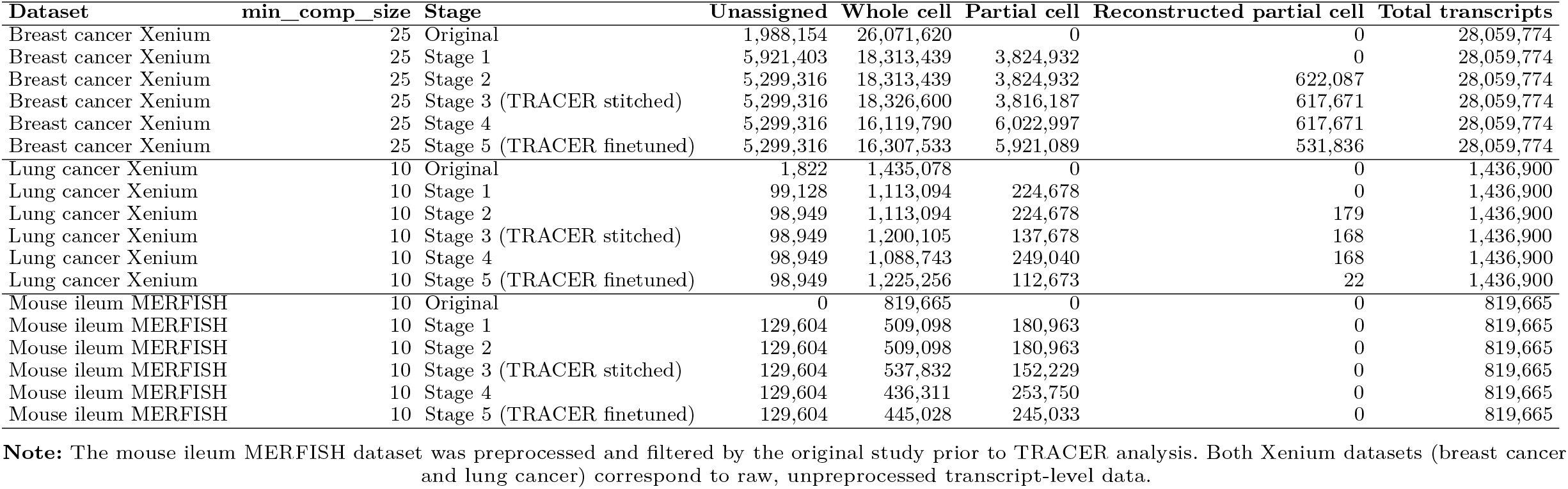
Transcript assignment across TRACER stages on three benchmark datasets. For each dataset and TRACER stage, the table reports the number of transcripts assigned to whole cells, partial cells, reconstructed partial cells, or remaining unassigned.

| Dataset | min_comp_size | Stage | Unassigned | Whole cell | Partial cell | Reconstructed partial cell | Total transcripts |
| --- | --- | --- | --- | --- | --- | --- | --- |
| Breast cancer Xenium | 25 | Original | 1,988,154 | 26,071,620 | 0 | 0 | 28,059,774 |
| Breast cancer Xenium | 25 | Stage 1 | 5,921,403 | 18,313,439 | 3,824,932 | 0 | 28,059,774 |
| Breast cancer Xenium | 25 | Stage 2 | 5,299,316 | 18,313,439 | 3,824,932 | 622,087 | 28,059,774 |
| Breast cancer Xenium | 25 | Stage 3 (TRACER stitched) | 5,299,316 | 18,326,600 | 3,816,187 | 617,671 | 28,059,774 |
| Breast cancer Xenium | 25 | Stage 4 | 5,299,316 | 16,119,790 | 6,022,997 | 617,671 | 28,059,774 |
| Breast cancer Xenium | 25 | Stage 5 (TRACER finetuned) | 5,299,316 | 16,307,533 | 5,921,089 | 531,836 | 28,059,774 |
| Lung cancer Xenium | 10 | Original | 1,822 | 1,435,078 | 0 | 0 | 1,436,900 |
| Lung cancer Xenium | 10 | Stage 1 | 99,128 | 1,113,094 | 224,678 | 0 | 1,436,900 |
| Lung cancer Xenium | 10 | Stage 2 | 98,949 | 1,113,094 | 224,678 | 179 | 1,436,900 |
| Lung cancer Xenium | 10 | Stage 3 (TRACER stitched) | 98,949 | 1,200,105 | 137,678 | 168 | 1,436,900 |
| Lung cancer Xenium | 10 | Stage 4 | 98,949 | 1,088,743 | 249,040 | 168 | 1,436,900 |
| Lung cancer Xenium | 10 | Stage 5 (TRACER finetuned) | 98,949 | 1,225,256 | 112,673 | 22 | 1,436,900 |
| Mouse ileum MERFISH | 10 | Original | 0 | 819,665 | 0 | 0 | 819,665 |
| Mouse ileum MERFISH | 10 | Stage 1 | 129,604 | 509,098 | 180,963 | 0 | 819,665 |
| Mouse ileum MERFISH | 10 | Stage 2 | 129,604 | 509,098 | 180,963 | 0 | 819,665 |
| Mouse ileum MERFISH | 10 | Stage 3 (TRACER stitched) | 129,604 | 537,832 | 152,229 | 0 | 819,665 |
| Mouse ileum MERFISH | 10 | Stage 4 | 129,604 | 436,311 | 253,750 | 0 | 819,665 |
| Mouse ileum MERFISH | 10 | Stage 5 (TRACER finetuned) | 129,604 | 445,028 | 245,033 | 0 | 819,665 |

**Note:** The mouse ileum MERFISH dataset was preprocessed and filtered by the original study prior to TRACER analysis. Both Xenium datasets (breast cancer and lung cancer) correspond to raw, unprocessed transcript-level data.

## 6 Methods

### 6.1 Data acquisition and preprocessing

All spatial transcriptomics data were obtained from previously published work, including data from Xenium (standard and multimodal configurations) and MERFISH platforms. The breast cancer Xenium dataset with nucleus expanded (NE) segmentation and associated cell-type annotations dataset were obtained from the original study [23]. The lung cancer Xenium dataset with multimodal segmentation was obtained from a previously published study [25]. We resegmented this dataset with NE segmentation using Xenium Ranger to obtained the dataset used in Fig. 3. Xenium transcripts were filtered using a quality value threshold of QV *>* 30 prior to analysis. MERFISH data were obtained from the Baysor study demonstrating three-dimensional cell segmentation using Cellpose-based priors [5]. From this dataset, transcripts with nuclear probability greater than 0.9, as reported by the Baysor output, were considered nuclear and used for downstream analysis.

All analyses were performed independently for each tissue section unless otherwise stated.

### 6.2 Translating normalized pointwise mutual information (NPMI) to spatial transcriptomics

#### 6.2.1 Conceptual framework

The coherence framework underlying this work is built on the translation of normalized pointwise mutual information (NPMI), originally developed in natural language processing, to spatial transcriptomics under a cell-as-context / gene-as-word paradigm.

In natural language processing, a document defines a unit of co-occurrence, and words are binary events indicating presence or absence within that document. Importantly, PMI and NPMI are defined over event occurrence, not word frequency.

This formulation is robust to document length variation and focuses on semantic co-occurrence patterns.

We adopted this framework directly for biological data by defining:

- Context (document): a segmented nucleus
- Word (token): a gene
- Event *i*: gene *i* is present in the nucleus
- Event *j*: gene *j* is present in the nucleus
- Event (*i, j*): both genes *i* and *j* are present in the same nucleus

This formulation explicitly avoids dependence on transcript counts, which are subject to technical noise and segmentation artifacts, and instead emphasizes co-expression programs across cells, cross-lineage incompatibilities, and doublet or merged-cell signatures.

#### 6.2.2 Nucleus-centered binary gene representation

As nucleus segmentation is less noisy than cell segmentation, the segmented nuclei were treated as context windows, and genes were considered present within a nucleus if and only if at least *N*_min_ transcripts were detected inside its boundary. In this study, we used *N*_min_ = 2 . This yielded a binary presence–absence matrix

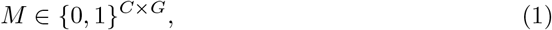

where *C* denotes the number of nuclei and *G* the number of genes.

No normalization, scaling, or transcript count weighting was applied beyond platform-specific quality filtering. This representation intentionally discards transcript abundance information in favor of robust detection of gene co-occurrence patterns across cells.

We selected nuclei rather than whole-cell segments as the primary context unit because nuclear segmentation is generally more reliable across platforms and tissues, whereas whole-cell boundaries are often inferred indirectly and may introduce systematic segmentation errors—particularly in standard Xenium datasets where cell outlines are extrapolated from nuclear signal. Using nuclei as contexts therefore provides a conservative and platform-agnostic basis for computing gene–gene associations.

#### 6.2.3 Empirical Probability Calculation

Let *C* denote the total number of nuclei. For each gene pair (*i, j*), we estimated marginal and joint probabilities of gene presence as

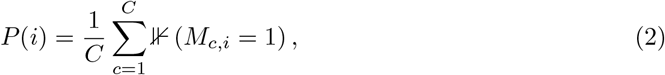

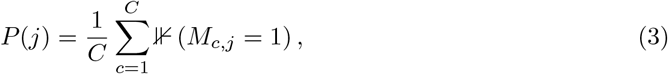

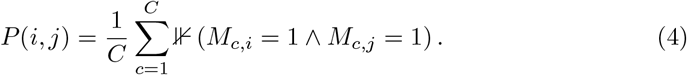

Each gene pair induces a complete four-part partition of the event space:

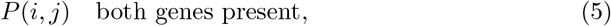

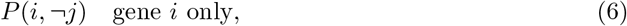

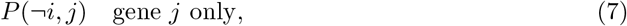

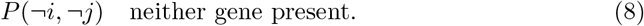

By construction, these probabilities sum to one for every gene pair, ensuring a valid probabilistic interpretation.

#### 6.2.4 Empirical NPMI computation

Pointwise mutual information (PMI) was empirically computed as

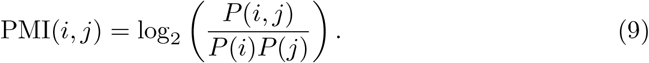

Because PMI is known to be biased toward rare events, particularly when *P* (*i, j*) is small, we used normalized PMI (NPMI):

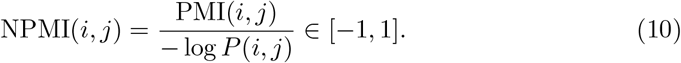

NPMI values of 0 indicate statistical independence, positive values indicate enrichment above independence, and negative values indicate depletion or mutual exclusivity. This normalization enables meaningful comparison across gene pairs with widely varying marginal frequencies.

The full gene–gene empirical NPMI matrix was computed in a reference-free, data-driven manner using nuclei selected to represent the central distribution of transcript counts for Xenium datasets (excluding the lowest and highest 20th percentiles) and all nuclei for MERFISH datasets.

### 6.3 Associative clique extraction and transcript consistency modeling

#### 6.3.1 Distinguishing whole cells and partial cells

In tissue-based spatial transcriptomics, sections of non-negligible thickness pass through cells at different cross-sections, potentially capturing cytoplasmic contents without the nucleus. Nuclear segmentation methods do not account for such non-nuclear profiles — when these occur in isolation, the cell is missed entirely; when they spatially overlap with a nuclear profile, their transcriptional contents are incorrectly attributed to that nucleus, introducing noise. To explicitly account for these possiblities, we distinguished between whole cells and partial cells based on combined imaging and transcriptomic criteria.

Whole cells were defined as segmented regions that exhibit a strong in-focus nuclear DAPI signal within the imaging plane, consistent with the presence of the nucleus and the majority of the cell body. In contrast, partial cells were defined as segmented regions containing cytoplasmic signal or transcripts but lacking a clearly in-plane nucleus, consistent with cells whose nuclei reside in adjacent sections along the *z*-axis. Partial cells arise frequently at tissue boundaries, within densely packed regions, or in areas with reduced nuclear signal, and may contribute spurious transcripts to neighboring cells.

Rather than discarding partial profiles, we treated them as explicit intermediate entities and incorporated them into downstream modeling in a conservative, coherence-driven manner.

#### 6.3.2 NPMI-based conservative transcript pruning

To resolve transcript assignment ambiguity while preserving biologically coherent gene programs, we implemented a two-pass conservative pruning strategy based on gene–gene support derived from NPMI.

For each segmented cell, the set of expressed genes was evaluated using the precomputed gene–gene NPMI matrix computed from all nuclei in the dataset. For a given cell, each gene was assessed based on its aggregate mutual support with other genes expressed within the same cell, defined by pairwise NPMI values.

In the first pruning pass, genes failing a predefined NPMI consistency criterion were removed from the original cell and reassigned to a corresponding partial-cell identifier. This step preserved the core, internally consistent gene set of each cell while isolating weakly supported or potentially spurious transcripts without discarding them.

In the second pruning pass, the same NPMI-based evaluation was applied to these partial gene sets. Genes that again failed the consistency criterion were conservatively reassigned as unassigned rather than being forced into any cellular entity.

This two-stage procedure minimized over-pruning while progressively separating transcriptionally coherent gene programs from inconsistent associations. Importantly, partial cells were retained as explicit entities, allowing the recovery of biologically meaningful but incomplete expression programs that would otherwise be lost.

### 6.4 Reconstruction of partial cells from unassigned transcripts

A subset of transcripts could not be confidently assigned to segmented cells, either because they were unassigned in the original segmentation or remained unassigned after two-pass pruning. Such unassigned transcripts may arise from extracellular regions that nonetheless reflect coherent cellular expression programs—for example, when a cell body lies within the analyzed plane but the nucleus is located in an adjacent section along the *z*-axis, or when nuclear signal is insufficient to pass automated segmentation thresholds.

To capture these structures, we introduced the concept of reconstructed partial cells, defined as spatially localized groups of transcripts lacking a prior segmentation mask but exhibiting strong internal spatial connectivity and transcriptional coherence consistent with a single cell.

Partial cell identification was performed using transcripts remaining unassigned after the two-pass pruning procedure, together with transcripts that were unassigned in the initial segmentation. These transcripts were embedded in three-dimensional physical space using their spatial coordinates and represented as nodes in a *k*-nearest neighbor (kNN) graph. Edges were formed between transcripts that were mutual nearest neighbors and lay within a fixed Euclidean distance threshold, enforcing local spatial connectivity. Connected components of this graph were extracted to yield candidate partial cell regions.

Components containing fewer than a minimum number of transcripts were discarded to reduce spurious aggregation. For the remaining components, transcriptional coherence was evaluated using the same NPMI framework applied to segmented cells. Within each component, genes were iteratively pruned using the aforementioned greedy NPMI-based strategy, removing transcripts whose genes lacked sufficient mutual support with the dominant gene co-expression program of the component. Components retaining internally consistent gene programs after pruning were labeled as partial cells, while transcripts removed during this step were conservatively discarded.

Partial cells were treated as distinct cellular entities in subsequent analyses and assigned unique identifiers, while maintaining explicit provenance from the unassigned transcript pool. This approach does not assume that unassigned transcripts are artifactual; instead, it enables recovery of transcriptionally coherent cellular profiles missed by nucleus-centered segmentation while preserving conservative handling of ambiguous signals.

### 6.5 Hierarchical stitching of cellular entities using Δ*C* coherence optimization

#### 6.5.1 Entity representation and spatial constraints

To reconcile fragmented cellular entities arising from segmentation artifacts and partial profiles, we performed constrained hierarchical stitching based on transcriptional coherence. Each entity—whole cell, partial cell, or reconstructed partial cell—was represented by its centroid in physical space and by the set of genes associated with its transcripts.

Candidate merges were restricted to spatially adjacent entities, defined using a Delaunay triangulation of entity centroids (computed in three dimensions unless otherwise stated). This constraint limited stitching to biologically plausible spatial neighborhoods.

#### 6.5.2 Coherence score based on gene–gene NPMI

For an entity with gene set *G* = {*g*_1_,…, *g*_*k*_}, we defined a coherence score

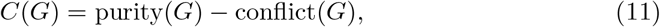

computed from the gene–gene NPMI matrix.

For all gene pairs (*g*_*i*_, *g*_*j*_) with *i < j*, and finite NPMI values *w*_*ij*_, purity and conflict were defined as

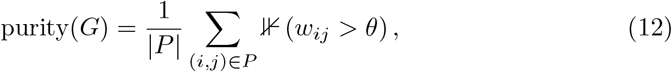

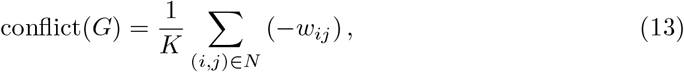

where *P* is the set of all observed gene pairs with finite NPMI, *N* ⊆ *P* denotes pairs with negative NPMI values, ⊮ (·) is the indicator function, *θ* = 0.05 accounts for biological noise, and 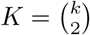 is the total number of possible gene pairs. If fewer than two genes were present or no valid gene pairs were observed, coherence was defined as zero.

Purity is normalized by the number of observed pairs |*P* | because it measures the fraction of gene pairs exhibiting strong positive co-expression. Conflict, in contrast, quantifies the total incompatible gene–gene association pressure, and is therefore normalized by the total number of possible gene pairs 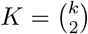. This choice ensures that conflict increases both with the number and magnitude of contradictory associations, providing a stable and size-consistent indicator of mixed-lineage profiles.

##### Mathematical properties of the purity and conflict score

To justify the statistical behavior and interpretability of the NPMI-derived purity and conflict functions, we establish several basic properties. Proofs are provided in Supplementary Note 1.

###### Remark 1

(Boundedness of purity and conflict)

Let *G* = {*g*_1_,…, *g*_*k*_} be a gene set expressed in a cell, let *w*_*ij*_ ∈ [−1, 1] denote gene– gene NPMI values, let *P* = {(*i, j*) : *w*_*ij*_ finite}, *N* = {(*i, j*) : *w*_*ij*_ *<* 0}, 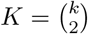 be the total possible gene pairs, and let *θ* ∈ [0, 1) be a small noise tolerance (e.g., 0.05). Then

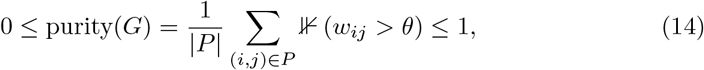

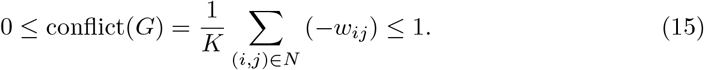

###### Remark 2

(Coherence score is bounded and well-posed)

Define *C*(*G*) = purity(*G*) − conflict(*G*). Then −1 ≤ *C*(*G*) ≤ 1.

###### Remark 3

***(Thresholding at*** *θ* ***removes stochastic noise under independence)***

Let *w*_*ij*_ be empirical NPMI estimated from Bernoulli co-occurrence in sparse spatial transcriptomics. Assume independence: *w*_*ij*_ ≈ 0 with sampling fluctuations on the order of 𝒪(*n*^−1*/*2^). Then

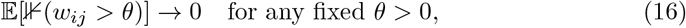

so 𝔼[purity(*G*)] ≈ 0 for independent gene pairs. The threshold *θ* is a noise floor ensuring that purity only reflects genuinely consistent positive associations, not stochastic sampling variance.

###### Remark 4

(Monotonicity)

Let *G* be fixed and consider a single gene pair (*a, b*). If *w*_*ab*_ increases and crosses the threshold *θ*, then purity(*G*) weakly increases and conflict(*G*) does not increase, therefore *C*(*G*) increases. If *w*_*ab*_ decreases further into the negative region, then conflict(*G*) strictly increases, purity(*G*) does not increase, and *C*(*G*) strictly decreases.

##### 6.5.3 ReLU-based coherence formulation

To reduce sensitivity to weak associations and emphasize stronger evidence, we additionally implemented a symmetric ReLU-based coherence formulation. For each gene–gene NPMI value *w*, we applied

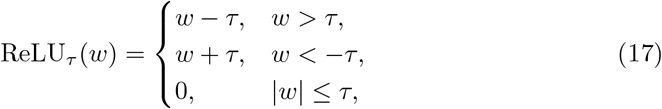

where *τ* defines a dead zone around zero. Similar to *θ*, we used *τ* = 0.05 to account for biological noise.

Purity and conflict were then defined as

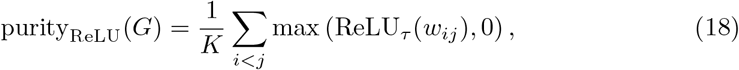

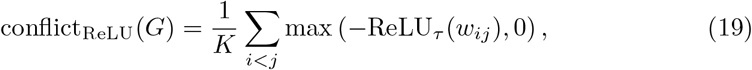

where 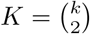. The coherence score was again defined as *C*_ReLU_(*G*) = purity_ReLU_(*G*)™ conflict_ReLU_(*G*).

###### Mathematical properties of the ReLU-based coherence formulation

To justify the statistical behavior and interpretability of the ReLU-based coherence formulation, we establish several basic properties. Proofs are provided in Supplementary Note 1.

###### Remark R1 (ReLU coherence preserves boundedness and monotonicity)

Since *w*_*ij*_ ∈ [−1, 1], the transformed values satisfy 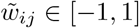. The ReLU transformation preserves sign ordering: increasing *w*_*ij*_ increases 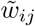, making *w*_*ij*_ more negative decreases 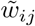. Therefore, 0 ≤ purity_ReLU_(*G*), conflict_ReLU_(*G*) ≤ 1, and *C*_ReLU_(*G*) remains bounded in [−1, 1]. Values with |*w*_*ij*_|≤ *τ* (dead zone) contribute exactly zero to both purity and conflict.

##### 6.5.4 Δ*C*-based merge criterion with simplicity penalty

For two candidate entities *U* and *V* with gene sets *G*_*U*_ and *G*_*V*_, we evaluated the change in coherence upon merging as

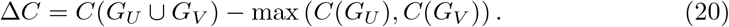

To discourage trivial merges driven solely by small gene sets, we optionally applied a simplicity penalty:

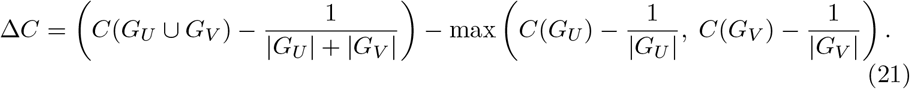

This penalty promoted absorption of smaller, fragmented entities into larger coherent ones while preventing over-merging of independent structures. Merges were accepted only when Δ*C* ≥ 0.

###### Mathematical properties of the ΔC merge criterion

To justify the statistical behavior and interpretability of the Δ*C*-based merge, we establish several basic properties. Proofs are provided in Supplementary Note 1.

###### Remark 5

(Δ*C* ***favors biologically coherent unions)***

Let *U* and *V* be two cell fragments. Then Δ*C* ≥ 0 only when the union does not introduce disproportionately many negative NPMI pairs. In particular, if both *G*_*U*_ and *G*_*V*_ are internally coherent but cross-pairs are conflicting, then Δ*C <* 0 and stitching is rejected; if cross-pairs reinforce the dominant gene program, Δ*C >* 0 and stitching is accepted.

##### 6.5.5 Constrained hierarchical stitching

Hierarchical merging was performed greedily using a union–find structure, prioritizing candidate merges with the largest positive Δ*C*. To preserve biological interpretability, merges between two entities that both contained a whole cell were disallowed.

After convergence, each stitched cluster was assigned a representative label, prioritizing whole cells, followed by partial cells, and then reconstructed partial cells. Spatial coherence was then enforced by splitting any stitched entity whose transcripts formed multiple disconnected spatial components, retaining the largest component under the original label.

As an optional post-processing step, transcripts remaining unassigned could be reassigned to the nearest partial cell, reconstructed partial cell, or optionally whole cell within a user-defined spatial distance threshold, enabling recovery of locally coherent transcripts without altering core assignments.

### 6.6 Coherence-derived metrics for segmentation evaluation

To enable quantitative evaluation of transcriptional coherence at the cellular level, we derived a set of metrics from the gene–gene NPMI framework. For a cellular entity with gene set *G*, purity and conflict scores were computed as defined above, using either the raw NPMI formulation or the ReLU-based formulation with dead-zone threshold *τ* .

We further defined the signal strength of a cellular entity as

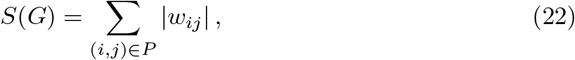

where *w*_*ij*_ denotes the NPMI value for gene pair (*g*_*i*_, *g*_*j*_). Signal strength captures the total magnitude of coherent and incoherent gene–gene associations within a cell.

To normalize coherence contributions, we defined relative purity and relative conflict as

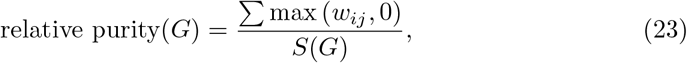

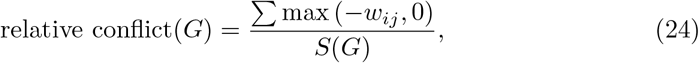

such that the two quantities sum to one when *S*(*G*) *>* 0 and at least one valid gene pair is present.

Transcript count per cellular entity was recorded as the total number of transcripts assigned to that entity after segmentation or refinement and used as a quality control measure. All metrics were computed independently for each cellular entity.

### 6.7 Single-cell analysis

Single-cell preprocessing and downstream analysis were performed using Scanpy [42]. For the breast cancer dataset, partial and reconstructed partial cells with fewer than 10 transcripts from at least 5 unique genes were removed; all other datasets included all cells. Raw counts were retained, followed by total-count normalization and log-transformation. PCA was computed on all genes to avoid variance-based gene selection bias. A *k*-nearest neighbor graph (*k* = 15) was constructed, and unsupervised clustering was performed using the Leiden algorithm (resolution = 1.0) with fixed parameters across datasets to ensure consistent interpretation of UMAP manifolds. UMAP embeddings were computed for visualization, and differential marker genes were displayed using Scanpy’s matrix plot function.

### 6.8 Scrublet-based expression doublet detection

Doublet detection was performed using the Scanpy wrapper for Scrublet [39] to benchmark segmentation-induced mixed-lineage profiles in an orthogonal expression-only framework. For each segmentation strategy (Baysor, TRACER-stitched, TRACER-fine-tuned), Scrublet was applied to the corresponding whole-cell expression matrix using default parameters to ensure comparability across methods. We evaluated three expected doublet rates (0.02, 0.2, 0.25, and 0.3) to assess robustness of doublet-score behavior across common parameter settings.

### 6.9 Ligand–receptor interaction analysis

Ligand–receptor (LR) interactions were inferred using Squidpy [34] with the OmniPath ligand–receptor database [35]. For each segmentation strategy (standard Xenium, TRACER-stitched, TRACER-fine-tuned), LR analysis was performed on the same AnnData object structure using identical parameters to ensure comparability.

Permutation-based testing was performed with *n*_perms_ = 100 and threshold = 0 to avoid filtering out ligands or receptors in sparsely expressed clusters. *P* -values were corrected for multiple testing using the Benjamini–Hochberg procedure, and interactions with false discovery rate (FDR) *<* 0.05 were considered significant.

### 6.10 Segmentation baselines and benchmarking details

For MERFISH mouse ileum, Proseg [36] was run in 3D mode with the following inputs: gene column `gene`, transcript ID column `mol_id`, coordinate columns `x`, `y`, `z`, prior cell-assignment column `cell`, unassigned value `0`, `–use-cell-initialization`, `–min-qv 0`, and `–nthreads 8`. Output was written as a SpatialData `zarr` object.

For Xenium breast cancer and lung cancer datasets, Proseg was run in Xenium mode with `–min-qv 30` and `–nthreads 8`.

For the lung cancer multimodal Xenium dataset, Baysor [5] v0.7.1 was run in 3D mode using transcript coordinates (`x`, `y`, `z`), gene identities (`feature_name`), minimum molecules per cell `-m 5`, polygon output enabled, and prior segmentation confidence `0.8`, using nucleus-expanded Xenium segmentation as the prior.

Standard Xenium breast cancer Baysor, Segger and BIDCell [37] outputs were obtained from the Segger study [38], and mouse ileum Cellpose and Cellpose+Baysor outputs were obtained from the Baysor study.

In the mouse ileum data, Proseg+Baysor consistently removed approximately 70% of transcripts across tested settings, including both 2D and 3D modes and more relaxed cell-size constraints.

### 6.11 Three-dimensional niche analysis with FunCN

We quantified local cellular neighborhoods using FunCN [33] on the TRACER-refined breast cancer Xenium dataset. We adapted FunCN to the three dimensions by replacing the original planar kernel-smoothing step implemented in the spatstat package with an explicit 3D Gaussian neighborhood model applied to cell centroid coordinates ((x,y,z)). For each target cell (i) and source cell (j), we computed a spatially weighted influence

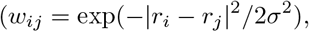

where *r*_*i*_ and *r*_*j*_ are the 3D coordinates of the *i*-th and *j*-the cell respectively. The kernel width *σ* was set to 30 *μ*m. We quantify the contribution of each cell type in the neighborhood of the target cell as the sum of the spatially weighted influences from all other cells in the tissue. To demonstrate the effect of TRACER on the neighborhood calculation, we first quantified the FunCN neighborhoods using only whole cells influences, and compared them to neighborhoods quantified using influence from using all TRACER-derived cellular entities, including partial cells.

### 6.12 Computational performance and scalability of TRACER

On a dual-socket Intel Xeon Platinum 8580 node (120 cores, 503 GiB RAM), TRACER processed a 28 million–transcript Xenium breast cancer dataset in approximately 5.2 hours using 17 CPU cores, with a peak memory footprint of 27 GiB (Supplementary Table 1). For smaller Xenium and MERFISH datasets (1.4 million and 0.8 million transcripts, respectively), end-to-end runtime was under 7 minutes and under 2 minutes. Across datasets, the most time-consuming components were the global coherence step (Stage 4) and final stitching (Stage 5), reflecting the cost of constructing and updating large spatial graphs.

## Declarations

### Data availability

All datasets analyzed in this study are publicly available. The multimodal Xenium lung cancer dataset (case: TSU-20, section: 1) can be downloaded from https://kero.hgc.jp/Ad-SpatialAnalysis_2024.html. The MERFISH mouse ileum dataset is available on Dryad (https://datadryad.org/dataset/doi:10.5061/dryad.jm63xsjb2). The standard Xenium breast cancer dataset can be accessed via https://www.10xgenomics.com/products/xenium-in-situ/preview-dataset-human-breast. The curated dataset used for benchmarking is available at Zenodo (DOI: 10.5281/zenodo.18892839)

### Code availability

The code for TRACER is publicly available in the GitHub repository https://github.com/imlong4real/TRACER under the Apache License 2.0.

## Acknowledgements

The authors would like to thank the Breakthrough Cancer Glioblastoma and Data Science TeamLab members, Alexander Ling, Nathalie Agar, Kenny Yu, Charles Couturier, Jennifer Gantchev, Wesley Tansey, and Christopher Tosh for their insightful feedback and assistance.

